# Environmental drivers of abundance and later seasonal activity of a salt marsh mosquito

**DOI:** 10.64898/2026.11.24.768890

**Authors:** Nadja C. Wipf, Thyge P. Enevoldson, Christopher M. Jones, Jennifer S. Lord, Tijani A. Sulaimon

## Abstract

1. Climate and environmental change are altering the seasonal dynamics of insects worldwide, yet long-term datasets remain scarce for mosquito vectors in temperate regions. Consequently, the environmental drivers and phenological responses of mosquito populations are still poorly resolved in northern Europe.

2. Here, we test whether the seasonal timing and abundance of a specialist coastal mosquito are jointly shaped by climatic and tidal drivers, and whether its activity period has extended over a 14-year monitoring record.

3. We analysed a 14-year dataset (2012–2025) of the salt marsh mosquito *Aedes detritus* – one of the most common saltmarsh mosquitoes in Europe - collected from four British coastal wetland sites. Weekly catch data from the full time series were used to assess long-term phenological trends, while generalised additive models were applied to site-month aggregated observations from the consistent four-site trapping period (2019– 2025) to identify climatic and tidal drivers of abundance.

4. Across the full 14-year record, the late-season peak and the end of the activity period (the 90% activity week) shifted later by 0.52 and 0.76 weeks per year, respectively (equivalent to fitted shifts of approximately 7 and 10 weeks), with no corresponding advance in the spring onset. This delayed autumn activity was robust to sampling-window and moving-window controls.

5. Adult abundance showed a pronounced bimodal seasonal pattern, with a smaller peak in late spring and a larger peak in mid-autumn. After accounting for seasonal structure, rainfall at a one-month lag and tidal surge were both positively associated with mosquito abundance. The size of the spring peak was correlated with rainfall, but the size and timing of the autumn peak were not.

6. These findings provide evidence that the seasonal activity of a salt marsh mosquito is extending later into autumn. We show that its population dynamics reflects a combination of climatic and tidal drivers whose relative contributions will require longer or more finely resolved records to disentangle fully. Sustained, long-term, full-season monitoring of coastal mosquito populations would therefore be important for understanding their responses to climate change. We recommend extending mosquito sampling in salt marsh areas into late autumn to better assess future risks associated with mosquito-borne pathogens.

## Introduction

Insects in coastal wetlands face environmental conditions that range from those that are relatively predictable and oscillatory such as the tide, to more unpredictable factors such as rainfall and turbulent coastal winds. Typically, coastal systems including salt marshes, are dominated by invertebrates which are habitat specialists rather than generalists (Rippel et al., 2021; Yang et al., 2009). Surviving within these habitats requires adaptations to acute and often daily fluctuations in salinity, the influx of predators and water depth, to name just a few examples. Populations of coastal insects are therefore shaped by the interplay between the climate and the tide (Rippel et al., 2021).

The effect of climate change on the life-history of insects is well documented (Chen et al., 2011; Wagner, 2020). One of the key conclusions is that climate change is influencing the timing of key behaviours and life events (the phenological response) (Bell et al., 2019; Bloomfield et al., 2026; Forrest, 2016; Guralnick et al., 2023). For example, spring advances of emergence or activity are commonly associated with warming temperatures (Bell et al., 2015; Roy et al., 2015), although responses can vary by taxa, habitat and latitude (Bell et al., 2019; Forrest, 2016). Most phenological studies focus on well-studied and charismatic species such as butterflies and moths for which long-term monitoring datasets are available (Bell et al., 2015). By contrast, the impact of climate change on species that occupy narrower ecological niches is, by comparison, poorly understood in temperate regions.

Salt marsh mosquitoes are highly adapted to brackish waters of coastal ecosystems across the world. These species (which mainly occur in the *Aedes* genus) lay their eggs in the moist soil of upper marsh land; requiring multiple inundations by saline tidal water to hatch. The timing and extent of tidal flooding therefore constrains when, and where, larvae are able to develop (Service, 1968) although rainfall is important in maintaining soil moisture and larval development (Yang et al., 2009), and as is the case for all mosquitoes, adult survival rates are influenced by several abiotic factors. Because climatic and tidal regimes vary over different timescales, disentangling their relative contributions to population abundance and phenology is difficult, but important to understand how coastal insects will respond to future environmental change.

The population structure of salt marsh mosquitoes can vary depending on the extensiveness of suitable brackish habitat and barriers to gene flow such as water crossings or vector control (Schmidt et al., 2025). Salt marsh species are highly dispersive (Jones et al., 2025), which may facilitate range expansion although the evidence for this remains equivocal. That said, due to their narrow niche, most populations are patchily distributed, and this limits their ability to expand their geographic range in response to climatic stress – a common response for many organisms (Hill et al., 2011; Schmidt et al., 2025).

Long-term and structured insect monitoring programmes are rare, yet they provide some of the strongest evidence for temporal change in insect populations and enhance our ability to detect the ecological drivers of that change (Shortall et al., 2024, 2009). Most insect monitoring data derive from snapshot surveys or contain considerable gaps within and among years, and so fail to capture the high year-to-year variation typical of insects (Cooke et al., 2025); this reduces power to detect trends and makes it difficult to separate genuine environmental associations from artefacts of inconsistent sampling (Didham et al., 2020). In temperate regions, there is growing concern over the climate-driven temporal and spatial expansion of mosquito activity and associated risk of pathogen (e.g. arboviral) transmission (Bruce et al., 2025; Tegar et al., 2026). The poleward spread of mosquito vectors in Northern Europe, for example, has received much attention, but comparatively fewer studies document inter-annual trends of temperate mosquito vector populations on a weekly basis, especially for over a decade or more.

Here, we use a structured, 14-year weekly dataset of the adult salt marsh mosquito, *Aedes detritus*, collected adjacent to the Dee Estuary salt marsh, Cheshire, UK to examine the environmental drivers of salt marsh mosquito abundance and decadal variation in length of active season. As *Ae. detritus* is highly adapted to the salt marsh and no other populations occur within the range of dispersal, this was an opportunity to investigate phenological change in a specialist, multivoltine species that is unlikely to be able to expand its range geographically, is a nuisance biter across Europe and an important potential public health risk due to its ability to carry multiple viruses pathogenic to birds and humans (Blagrove et al., 2016; Mackenzie-Impoinvil et al., 2015). Because the saltmarsh mosquito *Ae. detritus* breeds in high-marsh pools, we hypothesized that adult abundance would be positively associated with tidal conditions that inundate the high-marsh breeding pools and lagged rainfall, which maintains larval habitat. Furthermore, given regional warming trends, we hypothesized a long-term extension of the adult activity season. However, because *Ae. detritus* exhibits a bimodal annual cycle; we anticipated that early- and late-season phenology might change independently and show different environmental associations.

Specifically, we addressed four questions: (1) Which environmental variables best explain monthly adult abundance? (2) Does the bimodal seasonal cycle require separate early- and late-season phenology metrics? (3) Is there evidence for an extension of adult activity once uneven monitoring is accounted for? and (4) Are rainfall or cross-season carry-over patterns consistent with the phenology signals observed in the weekly data?

Our 14-year weekly trapping record is well suited to address these questions because – for the most part - it captures the smaller spring and larger autumn generation of *Ae. detritus*. This combination of weekly resolution, multi-year length and broad season coverage is rare among mosquito monitoring datasets and allows us to jointly estimate environmental drivers of abundance and long-term phenological change in a single specialist coastal species.

## Methods

### Study system

Of the 35 mosquito species established and native to the British Isles (Chandler, 2025), only a handful are associated with coastal wetlands. Hawkes et al. (2020) consider *Aedes detritus* and *Anopheles atroparvus* as the only truly brackish British species, with four further species – *Ae. caspius*, *Ae. dorsalis*, *Ae. flavescens* and *Culex modestus* – coastal-associated but not strictly saline breeders. *Aedes detritus* is the most abundant and widespread of these species, breeding in isolated brackish pools left by spring high tides or after heavy rains and emerging in large, synchronised broods (Hawkes et al., 2020; Service, 1968). Persisting in an environment few other mosquitoes can tolerate demands a distinct suite of adaptations – osmoregulation against saline water (Bradley et al., 1984), egg and larval development depending on sufficient rainfall or high enough tides to fill breeding pools – that set saltmarsh species apart from the freshwater-breeding mosquitoes most commonly studied. Adult dispersal ranges of up to 10 km in search of a host (Hawkes et al., 2020; Jones et al., 2025) and very occasionally *Ae. detritus* inland populations have been reported some 50 km from the coast (Marshall, 1933; Service, 1972). As a result, their phenology and population dynamics likely follow fundamentally different drivers than classical freshwater mosquito species.

### Mosquito trapping data

Weekly *Aedes detritus* abundance data were obtained from a longitudinal mosquito trapping study conducted at sites adjacent to the Dee Estuary, Cheshire, UK, from 2012–2025 (Figure 1) (Clarkson and Enevoldson, 2021; Clarkson and Setzkorn, 2011). At each site, adult mosquitoes were sampled using the Mosquito Magnet^®^ trap (Independence, Liberty Plus and Executive), baited with octenol (Octenol Finland). Records were kept of the number of mosquitoes collected and sampling effort quantified as the number of trap-days.

**Figure 1.**
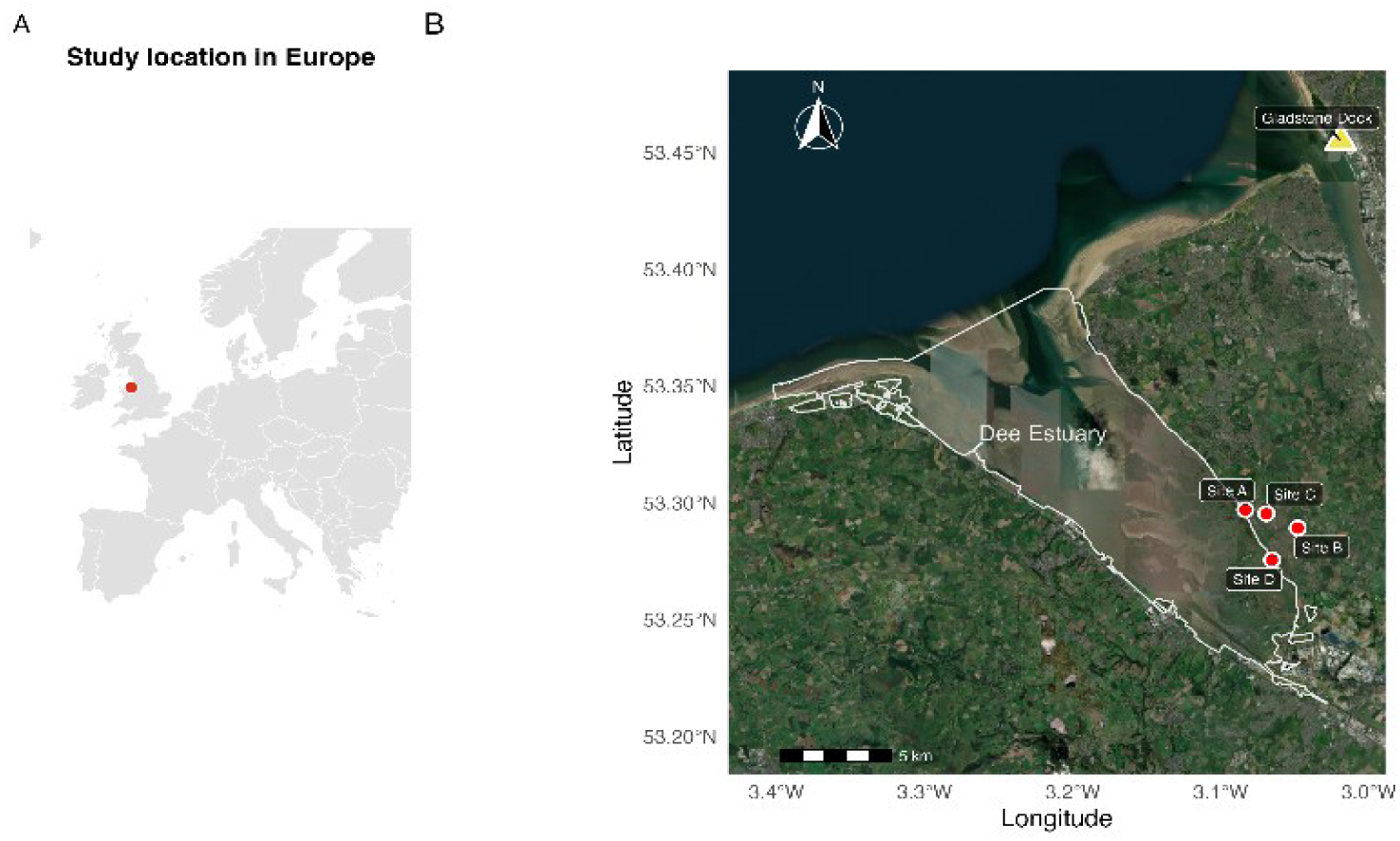
Study area and mosquito trap locations on the Dee Estuary, UK. Satellite imagery showing the interface between residential areas and the salt marsh. Red points indicate the approximate locations of the four long-term trapping sites (A–D). The yellow triangle marks the Gladstone Dock at the Port of Liverpool from where tidal data were derived. Base satellite imagery was generated in R using the *maptiles* package (Giraud et al., 2025).

From 2019 onward, traps were operated at four fixed sites with consistent seasonal coverage (Figure S1). We used this consistent four-site trapping period (2019–2025) to analyse the environmental correlates of monthly abundance, aggregating weekly observations to the site– month level. Data from all available sites and years (2012–2025) were retained for the phenology analyses, for which catches and trap-days were pooled with calendar week.

### Climate and environmental data

For the period from January 2012 to December 2025, we obtained monthly rasterised climate data from the Centre for Environmental Data Analysis (CEDA) (Met Office et al., 2026) and tidal data for Liverpool-Gladstone Dock from the British Oceanographic Data Centre (BODC, available from https://www.bodc.ac.uk/data/hosted_data_systems/sea_level/uk_tide_gauge_network/processed/; Open Government Licence v3.0). Climate variables included mean air temperature (°C), total precipitation (mm), mean relative humidity (%), and mean 10-m wind speed (m/s). Tidal variables included the monthly mean sea level (mean tide in m), the minimum and maximum observed sea level during the month (extremes), and the monthly minimum and maximum of the measured-minus-predicted sea-level residual, hereafter designated minimum and maximum surge.

Climate rasters were processed in R using the *terra* package (Hijmans et al., 2026). Values were extracted at each trap coordinate using bilinear interpolation to produce site-specific monthly time series. Tidal data were derived from a single station (Gladstone Dock, Liverpool, UK; Figure 1) and assumed representative of the study area. Missing tidal observations were imputed by Kalman smoothing of a structural time-series model implemented in *imputeTS* (Moritz et al., 2025; Figure S2). Before imputation, we recorded whether each monthly tidal value was present in the original BODC data. These records were kept after imputation, allowing the candidate models to be refitted in a sensitivity analysis using only site-months for which minimum surge (the tidal variable in the selected model) had been directly observed.

To account for delayed climatic effects on larval development and adult emergence, temperature and rainfall were lagged by one month. Lags were calculated on the complete calendar-month environmental series before catch data were joined, preventing winter gaps in trapping from linking the first sampled month of one year to the preceding autumn. Wind speed, relative humidity and tidal variables were modelled without a lag. Continuous environmental variables were standardised to a mean of zero and a standard deviation of one.

### Statistical analysis: Drivers of monthly abundance

To investigate environmental associations with monthly mosquito abundance, we fitted generalized additive models (GAMs; *mgcv* package in *R*, (Wood, 2025, 2017)) with a negative binomial error distribution and log link function to account for substantial overdispersion in the mosquito count data. We selected GAMs due to their flexibility in capturing non-linear relationships between abundance and environmental variables. The total monthly catch at each site was the response variable, with the natural logarithm of monthly trap-days included as an offset to account for variation in sampling effort.

Lagged temperature and same-month relative humidity were strongly seasonal, with calendar month explaining 94% and 69% of their variance, respectively (Figure S3). Including these variables alongside the month smooth term produced high concurvity; this made their separate effects difficult to interpret (Wood, 2017). One-month lagged rainfall, wind and the tidal summaries showed weaker seasonal associations. We therefore compared two representations of seasonal structure. Hypothesis H1 used a smooth of calendar month, which represented the combined effects of temperature, photoperiod, humidity and other correlated seasonal factors. Hypothesis H2 replaced month with explicit smooths of one-month lagged temperature and same-month relative humidity. Within each hypothesis, candidate models added one-month lagged rainfall, one of two alternative tidal terms (minimum surge or maximum extreme tide), and wind speed. No model contained more than one tidal summary. Site-plus-year and effort-offset-only null models provided baselines for model comparison.

Except for the effort-offset-only null model, all candidate models included a site-specific random intercept, year as a categorical variable and the log trap-effort offset. Year was modelled as a factor because the seven-year series was too short to estimate a reliable long-term smooth and showed substantial non-monotonic variation. Environmental predictors were fitted as thin-plate regression splines with *k*=4, except temperature and humidity, for which *k*=5 was required by the basis-dimension checks. Calendar month was fitted using a non-cyclic thin-plate spline with *k*=10 because trapping did not cover a complete annual cycle. Models were fitted by restricted maximum likelihood and ranked using Akaike’s Information Criteria corrected for small sample sizes (AICc) (Burnham and Anderson, 2002) implemented in the *MuMIn* package (Bartoń, 2026). Akaike weights quantified relative support for each model within the candidate set.

The selected model was checked for concurvity, basis-dimension adequacy, simulation-based residual patterns using the *DHARMa* package (Hartig et al., 2024), and agreement between observed and fitted values. To ensure our findings were robust to anomalous data and imputation, the candidate set was refitted after excluding the exceptionally high-abundance year of 2023 and separately refitted using only site-months with directly observed minimum surge.

### Analysis of seasonal phenology

#### Phenology metrics

Catches and trap-days were summed across all sites within each year and calendar week, and weekly activity was expressed as catch per trap-day. Season start and season end were the first and last sampled weeks with a positive catch, respectively, but were treated as boundary-sensitive because a single mosquito could change either metric. The 10%, 50% and 90% activity weeks represent the first calendar weeks at which the cumulative sum of weekly catch rates reached those proportions of the annual summed catch rate.

The long-term mean seasonal activity showed a mid-season trough around week 29 (Figure S4). We therefore divided each year at calendar week 30, defining an early and late season window before and after week 30 respectively. Within each window, peak timing was the week with the highest catch rate and peak magnitude was the corresponding maximum weekly catch per trap-day. A single global peak week was calculated only for comparison because changes in the relative magnitudes of the early- and late-season peaks could cause it to switch between spring and autumn.

#### Phenological trends and sampling-window controls

Annual phenology metrics were regressed by year using ordinary least squares, with slopes reported in weeks per year together with 95% confidence intervals and *p*-values. Trends were estimated for the consistently sampled period (2019-2025), the full series (2012-2025), and the full series excluding 2017 and 2018. The latter two years were identified as possible confounders due to inadequate sampling coverage: sampling only began at week 25 in 2017 and ended at week 37 in 2018 while mosquitoes were still being caught (Figure S5).

To assess the sampling-window biases (Didham et al., 2020), we used three sensitivity checks. First, metrics were recalculated within a common boundary window defined by the latest initial sampling week and the earliest final sampling week across all years. Second, trends were fitted across every contiguous period of at least four years. Third, we excluded 2017 data when evaluating early onset metrics and 2018 when evaluating late-season metrics, according to which seasonal boundary was censored in each year.

#### Environmental correlates and cross-season carry-over

Spearman rank correlations were used to explore associations between annual phenology metrics and the environmental variables included in the selected abundance model (rainfall and minimum surge). Monthly climate values were averaged across sites; rainfall was summed and minimum surge averaged within each defined seasonal window. Correlations were calculated for the full series, the consistent 2019–2025 period, and the series excluding 2017 and 2018. Spring phenology was tested against concurrent (March–May) and lagged (February–April) rainfall, with shorter windows (March–April, April) examined post hoc. Late-peak metrics were compared against July–September conditions, and the 90% activity week against August–October conditions. To account for multiple comparisons, *p*-values for all 24 correlations within each sampling period were adjusted using the Benjamini–Hochberg procedure (reported as *q*-values).

Motivated by Macgregor et al. (2019), we tested whether the later activity in year *t* was associated with a larger early-season peak in year *t* + 1, using only consecutive year pairs. The 90% activity week and central 80% activity span were tested in the raw data, after excluding 2017 and 2018, and after applying the common-window control.

## Results

### Seasonal activity of Aedes detritus was bimodal, with more consistent late-season peaks

Across all sites and years, the dataset contained 38,506 *Ae. detritus* from 1,185 trapping records and 2,573 trap-days. The four-site subset used for the abundance models contained 33,864 mosquitoes from 923 trapping records and 1,831 trap-days, which aggregated to 229 site-month observations. Sampling effort became substantially more regular from 2019 (Figure S1), although the first and last sampled weeks and the total number of sampled weeks continued to vary among years (Figure S5).

Weekly catch rates showed two distinct activity periods separated by a mid-season trough near week 30 (Figure 2; Figure S6). Early-season peaks occurred in some years but were weak or absent in others, whereas late-season activity occurred more consistently. The highest catch rates were recorded in 2023 and 2024.

**Figure 2.**
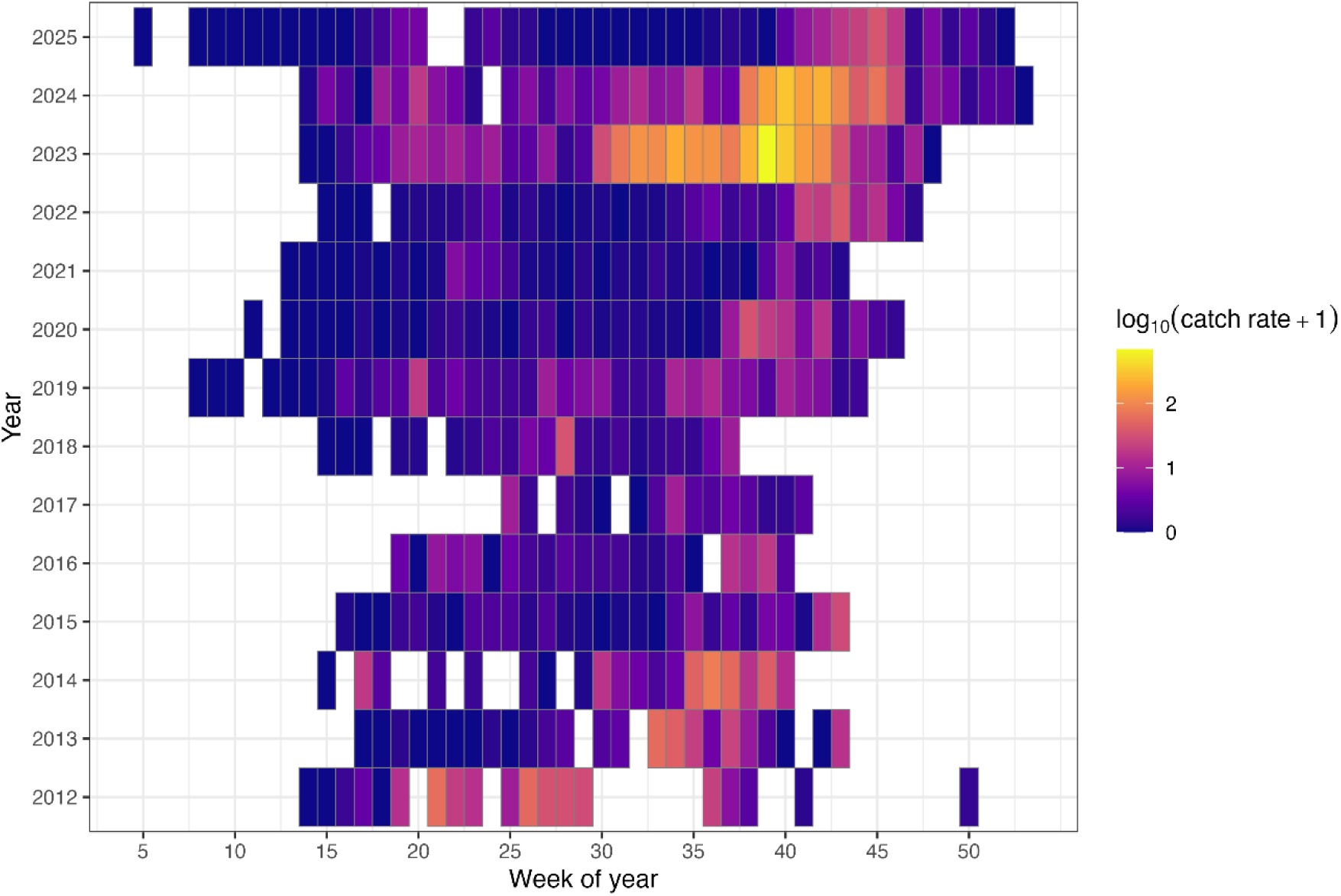
Heatmap showing weekly *Aedes detritus* catch-rate pooled across all trapping sites, 2012–2025. Coloured cells represent sampled weeks showing log10(catch per trap-day + 1) with yellow indicating high and blue low catch rates. White cells represent unsampled weeks. Weeks are consecutive seven-day bins from 1 January.

### Seasonality, lagged rainfall and minimum surge best explained monthly abundance

Monthly abundance followed a bimodal seasonal cycle, with a smaller rise in late spring and a larger rise in mid-autumn (Figure 3). The top-ranked model comprised a calendar-month smooth, one-month lagged rainfall and minimum surge (S + R + Su; df = 20, AICc = 1781.4, Akaike weight = 0.62; Table 1). The second-ranked model contained the same terms plus wind speed (S + R + Su + W; ΔAICc = 1.0, weight = 0.37). We therefore used the more parsimonious top-ranked model for effect estimation and interpreted the evidence for an additional wind effect as uncertain.

**Figure 3.**
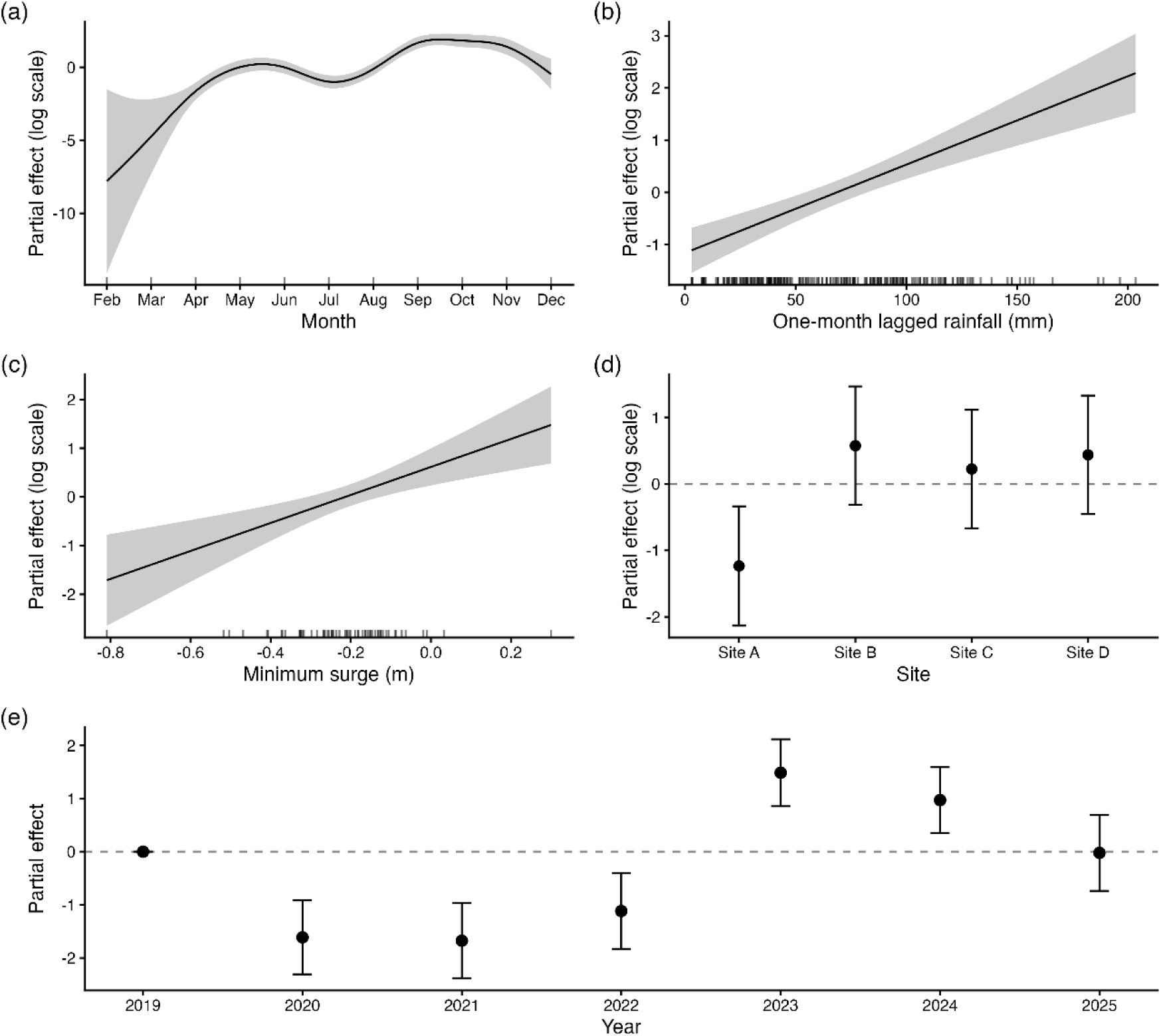
Partial effects from the selected monthly negative-binomial GAM: (a) calendar month, (b) one-month lagged rainfall, (c) minimum surge, (d) site and (e) year. Smooth lines and shaded bands show estimated partial effects and 95% confidence intervals; points and error bars show estimated site and year effects with 95% confidence intervals. Effects are on the log scale, and rug marks show the distribution of observations.

**Table 1.** Complete negative-binomial generalized additive models (GAM) candidate set to investigate environmental drivers of monthly mosquito abundance from 2019 to 2025.

| **Hypothesis** | **Model** | **df** | **AICc** | **ΔAICc** | **weight** |
|---|---|---|---|---|---|
| H1 | S + R + Su | 20 | 1781.4 | 0.0 | 0.62 |
| H1 | S + R + Su + W | 23 | 1782.4 | 1.0 | 0.37 |
| H1 | S + R | 19 | 1791.3 | 9.9 | 0 |
| H1 | S + R + W | 22 | 1791.6 | 10.3 | 0 |
| H1 | S + R + Xt | 20 | 1793.4 | 12.1 | 0 |
| H1 | S + R + Xt + W | 23 | 1794.2 | 12.8 | 0 |
| H2 | T + H + R + Su | 20 | 1801.3 | 19.9 | 0 |
| H2 | T + H + R + Su + W | 21 | 1803.2 | 21.8 | 0 |
| H2 | T + H + R | 19 | 1816.1 | 34.8 | 0 |
| H2 | T + H + R + Xt | 22 | 1817.3 | 36.0 | 0 |
| H2 | T + H + R + W | 20 | 1818.3 | 37.0 | 0 |
| H2 | T + H + R + Xt + W | 23 | 1819.0 | 37.6 | 0 |
| H1 | S | 17 | 1824.7 | 43.3 | 0 |
| H2 | T + H | 18 | 1838.3 | 57.0 | 0 |
| Null | Null (site + year) | 10 | 1955.9 | 174.5 | 0 |
| Null | Null (intercept) | 2 | 2081.5 | 300.1 | 0 |

*Note*: Models are ranked by small-sample corrected Akaike Information Criterion (AICc). ΔAICc indicates the difference from the best model, and Akaike weights represent the relative probability that a given model is the best in the candidate set. S = calendar-month smooth (seasonality), R = one-month lagged rainfall (mm), Su = minimum surge (m), W = wind speed (m/s), Xt = maximum extreme tide (m), T = air temperature (°C), and H = relative humidity (%). H1 are hypotheses with integrated seasonality; H2 with explicit environmental drivers; Null the null hypotheses. All models include site random effects, year as categorical factor, and a log trap-effort offset.

Models without minimum surge were at least 9.9 AICc units behind the top-ranked model, supporting the contribution of minimum surge. The best H2 model, which represented seasonality using lagged temperature and same-month relative humidity rather than calendar month, was 19.9 AICc units behind the top-ranked model. This indicated that calendar month represented seasonal structure more effectively than the explicit climate terms. The site-plus-year and effort-offset-only null models were 174.5 and 300.1 AICc units behind, respectively. Their poor support showed that trapping effort, site and year alone did not account for the observed monthly variation.

The selected model explained 74.1% of the deviance. Calendar month had a non-linear association with abundance (edf = 6.78, *p* < 0.001), describing an early-season rise, a mid-season trough and a larger late-season rise (Figure 3). Lagged rainfall and minimum surge had approximately linear effects (edf = 1.00). Abundance increased with lagged rainfall and as monthly minimum surge became less negative. The corresponding χ^2^ statistics were 37.27 and 14.11, and both terms had *p* < 0.001 (Figure 3; Table S1). Site effects differed significantly (edf = 2.88, *p* < 0.001), while year effects were largest in 2023 and 2024.

When the candidate models were refitted without the exceptionally high-abundance year 2023, the wind-inclusive model (S + R + Su + W) ranked first, with an Akaike weight of 0.90 (Table S2). When they were refitted using only site-months with directly observed minimum surge, S + R + Su ranked first, although the wind-inclusive model received similar support, with ΔAICc = 0.3 (Table S3). Lagged rainfall and minimum surge were therefore in the top-ranked model in both analyses. In-sample fitted values captured the broad seasonal pattern at all four sites, although some peak magnitudes were under- or overestimated, particularly in 2023 and 2024 (Figure S7). Simulation-based residual diagnostics found no significant evidence of non-uniformity, dispersion problems or excess outliers (Figure S8).

### Extension of late-season activity

The relative sizes of the early- and late-season catch peaks varied among years, so the global annual maximum sometimes switched between them (Figure S6). We therefore analysed early- and late-season peaks separately, and used cumulative activity weeks rather than first and last positive catches to reduce the influence of isolated early and late catches. Because cumulative metrics can still be affected by missing early- or late-season sampling, we then tested whether the trends changed when we restricted the sampling weeks or excluded years with incomplete seasonal coverage.

Across the full series, the late-season peak moved later by 0.52 weeks per year (95% CI 0.11 to 0.93; *p* = 0.018), and the 90% activity week moved later by 0.76 weeks per year (95% CI 0.34 to 1.18; *p* = 0.002; Table S4; Figure S9). Statistical support was weaker over the shorter consistent-sampling period: the late-season peak moved later by 1.07 weeks per year (*p* = 0.045), while the 90% activity week showed a similar direction but did not meet the nominal significance threshold (+0.79 weeks per year, *p* = 0.056). Season start and early-peak timing showed no directional trends across the full series (-0.15 and -0.14 weeks per year; *p* = 0.417 and *p* = 0.553; Table S4). The 10% activity week moved later in the untruncated full series (+0.96 weeks per year, *p* = 0.038), but changed direction under the common-window controls and therefore did not provide consistent evidence of a later onset (Table S5). We next tested whether differences in annual sampling windows could explain the late-season trends.

### Late-season shifts persisted under metric-specific sampling-window controls

Only calendar weeks 25-37 were sampled in every year from 2012 to 2025. After restricting all annual series to these weeks, neither the late-season peak timing nor the 90% activity week was associated with year (Table S6). This interval excluded much of both early- and late-season activity, so it provided only a restricted view of the biological season.

After excluding 2017, onset metrics were recalculated within weeks 19-37; none was associated with year (Table S7). After excluding 2018, however, and including weeks 25 to 40, both the late-season peak (+0.39 weeks per year, *p* = 0.006) and the 90% activity week (+0.24 weeks per year, *p* = 0.005) were delayed (Table S8). Metric-specific controls therefore supported later late-season activity rather than an advance in onset or a uniform shift in both seasonal boundaries. The main comparison is summarised in the robustness table (Table 2), with results for all phenology metrics provided separately (Table S5).

**Table 2.** Robustness of the three phenology metrics used for inference. Raw and fixed-window cells are OLS slopes in weeks per year; an asterisk denotes nominal *p* < 0.05. Fixed-window cells also report the shared calendar weeks. The moving-window column gives the number of significant slopes among all 66 contiguous periods of at least four years, with the direction of those slopes in parentheses.

| **Metric** | **2019-2025** | **2012-2025** | **2012-2025**<br>**excl. 17/18** | **Fixed all** | **Fixed**<br>**excl. 17/18** | **Moving**<br>**windows** |
|---|---|---|---|---|---|---|
| Early-season peak | -0.57 | -0.14 | -0.11 | -0.03<br>(wk 25-37) | -0.30<br>(wk 19-40) | 3/66 (+/-) |
| Late-season peak | +1.07* | +0.52* | +0.49* | +0.03<br>(wk 25-37) | +0.38*<br>(wk 19-40) | 12/66 (+) |
| 90% activity week | +0.79 | +0.76* | +0.74* | -0.06<br>(wk 25-37) | +0.43*<br>(wk 19-40) | 19/66 (+) |

The moving-window analysis tested whether these conclusions depended on choosing 2012 or 2025 as the limits of the series (Figure 4). Early-season peak slopes varied around zero, whereas all 12 nominally significant late-season peak slopes and all 19 nominally significant slopes for the 90% activity week were positive. No season-start slope was nominally significant (Table 2). The complete metric set and numerical summary are provided in Figure S10 and Table S9.

**Figure 4.**
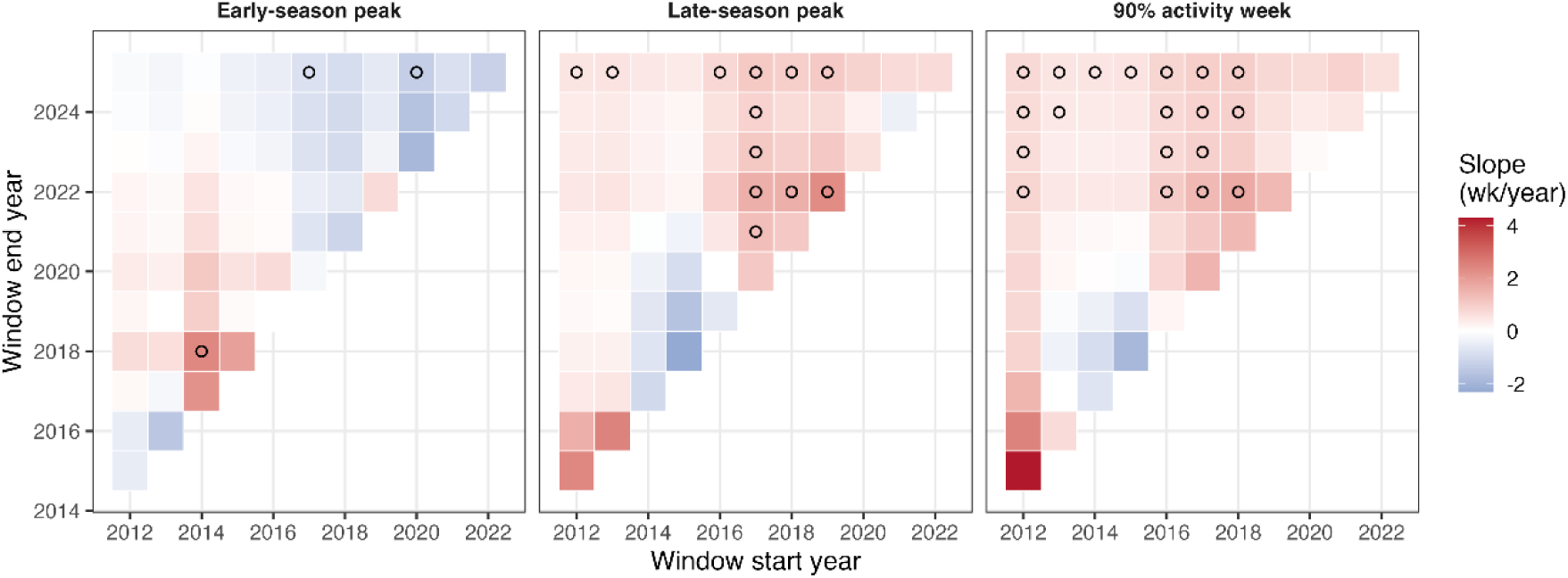
Moving-window trends for the three phenology metrics used for inference. Each tile is an OLS trend fitted to one contiguous start-end year period of at least four years. Red indicates later activity, blue earlier activity, and open circles denote nominal *p* < 0.05. Overlapping windows are descriptive stability checks rather than independent tests.

### Spring rainfall was associated with early-peak magnitude, but not the timing of the peak

Based on the evidence for a seasonal shift in the autumn activity of *Ae. detritus*, it is plausible that this could have a positive influence on the size of the following generation in the spring (Macgregor et al., 2019). We examined two aspects of this relationship: whether the magnitude of the spring peak was associated with rainfall, and whether the timing of late-season activity was associated with the size of the following spring peak.

Early-season peak magnitude was positively associated with spring rainfall, whereas peak timing was not (Figure 5). Across the full series, the association was stronger for rainfall during March-May (*ρ* = 0.727, *p* = 0.003) than during February-April (*ρ* = 0.464, *p* = 0.095; Table S10). The same contrast was seen during the consistent period and after excluding 2017 and 2018 (Table S11; Table S12).

**Figure 5.**
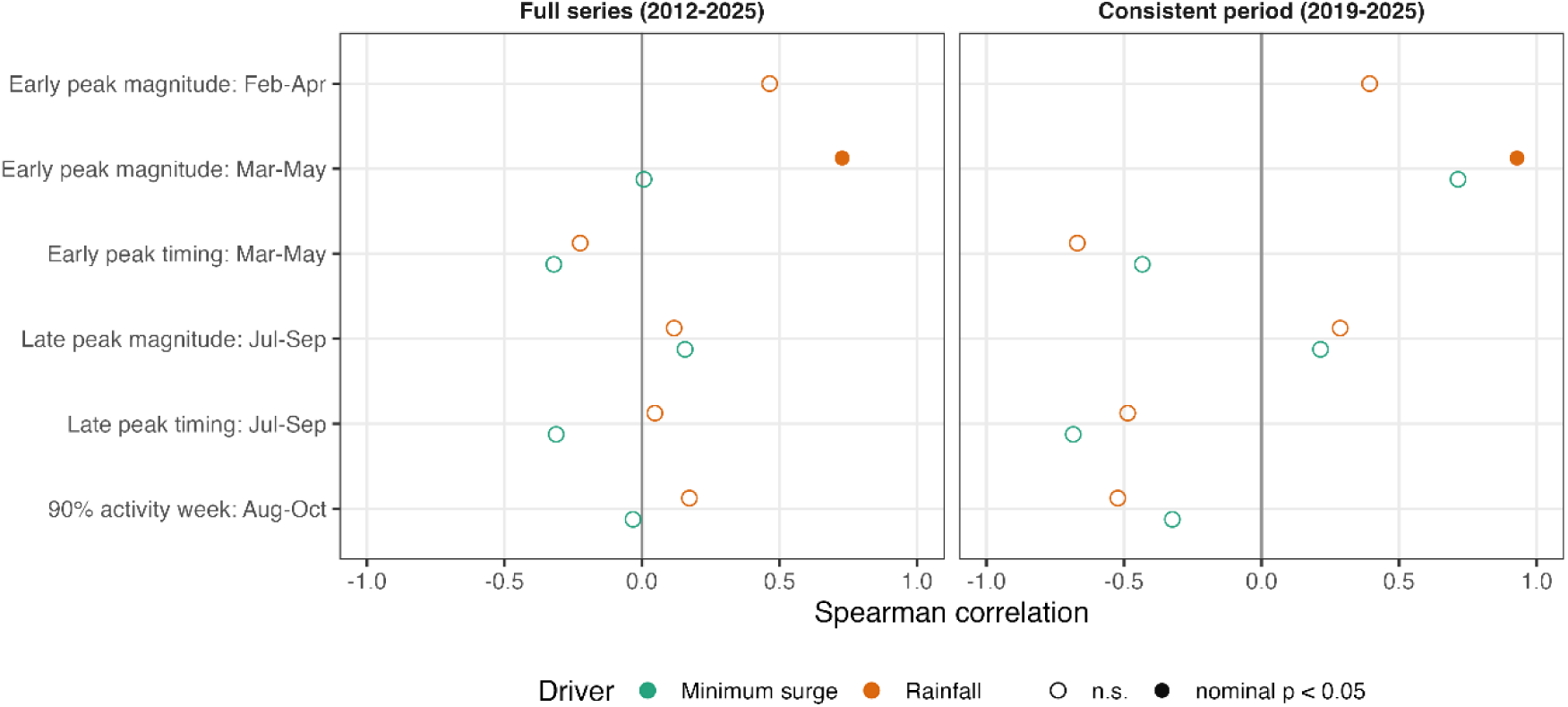
Focused annual associations between the abundance-model drivers and phenology metrics. Points show Spearman correlations for the full series and the consistent sampling period. Orange denotes rainfall and green minimum surge; filled points have nominal *p* < 0.05. Multiple-testing-adjusted *q*-values were calculated across all 24 tests within each period and are reported in the text and supplementary tables. Complete correlations for all phenology metrics, drivers and sampling periods are shown in Figure S11.

The narrower March-April and April-only windows also yielded positive associations (Figure S11). All four spring windows included April, so the comparisons could not determine whether rainfall acted before emergence or during spring activity. After adjustment across 24 tests per period, no association met *q* < 0.05 (minimum *q* = 0.060).

Unlike the monthly abundance results, annual late-season timing was not associated with rainfall or minimum surge during the full or consistent periods (Figure 5). This conclusion was unchanged after excluding 2017 and 2018 (Table S12).

### Evidence for cross-season carry-over depended on sampling controls

A positive cross-season association appeared only after excluding 2017 and 2018 and applying the common-window control: later 90% activity weeks in year *t* were followed by larger early-season peaks in year *t* + 1 (*ρ* = 0.781, *p* = 0.008, n = 10; Figure S12). Neither the raw data nor exclusion of 2017 and 2018 alone supported this association, and activity span did not show the same pattern (Table S13).

## Discussion

We set out to test whether the seasonal timing and abundance of a specialist coastal mosquito are jointly shaped by climatic and tidal drivers, and whether its activity period has extended over 14 years of monitoring. We found that monthly *Aedes detritus* abundance was best explained by seasonal structure together with rainfall and tidal surge, and that its pronounced bimodal life cycle required separate early- and late-season phenology metrics. Seasonal activity did shift over the study period, expressed mainly as a later end to activity in autumn rather than an earlier start in spring, and this shift proved robust to sampling-window controls.

This later-autumn extension contrasts with the springtime advances that dominate reports of climate-driven phenological change in temperate insects (Bell et al., 2015; Bloomfield et al., 2026). Longer and warmer seasons can allow multivoltine species to fit in extra reproductive cycles and, in some cases, expand their geographic range polewards (Macgregor et al., 2019). *Aedes detritus* is confined to a narrow, patchily distributed high-marsh habitat, which make the kind of range expansion documented in other multivoltine insects less available to it than to habitat generalists. In such a case, our data show that one phenological response to a changing climate is the delay of activity period rather than an earlier start to the season. A delayed response has been demonstrated in other late-season temperate species (e.g. mayfly (Glazaczow et al., 2016)). For hematophagous insects, shifts in the first and last recorded *Culicoides* biting midges over 40 years in northern England were associated with abundance, temperature and rainfall (Sanders et al., 2019). *Aedes detritus* has several overlapping generations, but its combination of tidal and rainfall-driven habitat dynamics (see below) means that the environmental drivers of its phenological extension are likely to differ from those documented in freshwater or terrestrial systems.

Phenological metrics are dependent on the timing of sampling. Uneven monitoring windows or sticking to the same sampling calendar dates despite changes in seasonal insect activity, can bias trend estimates in either direction, an issue described as a ’groundhog effect’ (Didham et al., 2020). We addressed this by comparing raw trends against trends recalculated within a common sampling window and across moving four-year windows (Figure 4). Late-season peak timing and the 90% activity week showed a consistent later shift across most of these controls, whereas trends in early-season peak timing did not reach significance under any control. We are therefore confident in the extension of activity into autumn than in any shift in spring emergence, and we report the latter as a weaker, non-significant pattern rather than a confirmed trend. Because early season catches of *Ae. detritus* are typically smaller than the dominant autumn peak, this may partly reflect lower statistical power at the start of the season rather than a true absence of change.

The top-ranked model for monthly *Ae. detritus* abundance included a smooth seasonal term, one-month lagged rainfall and minimum tidal surge. Across the four sites, abundance followed a bimodal seasonal pattern, with a smaller rise in late spring followed by a larger rise in mid-autumn. This bimodal activity has also been observed elsewhere in UK wetlands (Hawkes et al., 2020). Lagged rainfall had an approximately linear positive association with abundance, consistent with recent findings from a shorter study in Scotland (Kirby et al., 2026). Minimum surge added information beyond seasonality and rainfall, with higher (less negative) values associated with greater abundance. Monthly summaries may nevertheless miss short-lived tidal inundation events so we interpret this with caution. The wind-inclusive model had similar support in the full dataset and became top-ranked after excluding 2023, so evidence for the effect of wind was less stable and may require finer-resolution analysis.

To some extent these findings confound our initial expectations, as we anticipated tidal patterns, rather than rainfall, to be more influential on *Ae. detritus* abundance. Salt marsh mosquitoes occupy high salt marsh habitat above mean high-water levels and repeated inundations of marsh pools with saline tidal water is needed for eggs to hatch into immature larvae(Service, 1968). As thetideretreats, eggs arelaidwithinexposedmoistsoilandcan remain dormant for several months until the next high tide. Density-dependent modelling on a 15-year salt marsh mosquito (*Aedes vigilax*) dataset in northern Australia, has shown that low tide events are predictors of mosquito population booms, with growth rates peaking two months prior to the eventual adult peak following the first high tide, supplemented with low rainfall (Yang et al., 2009). *Aedes detritus* egg hatching and larval development can occur across a wide range of salinities, with rainwater acting as a triggering factor (Clarkson and Setzkorn, 2011; Marshall, 1938; Service, 1972). This most likely explains the positive association between lagged rainfall and *Ae. detritus* abundance. That said, we caution that the monthly resolution of our tidal and rainfall covariates is coarse relative to the event-driven dynamics of salt marsh flooding and may not fully capture short-lived inundation events. Therefore, a finer-resolution analysis would be needed to establish whether monthly aggregation underestimates the contribution of the tide relative to that of other potential environmental drivers.

Although lagged rainfall and minimum surge were the strongest predictors of monthly abundance, neither variable was significantly associated with the year-to-year timing of the late-season shift and we could not attribute the observed extension of activity directly to either driver. This may reflect a lack of statistical power given only 14 data points (years) but may also point to additional mechanisms driving the phenological change in the autumn, such as longer average adult survival which may be affected by different variables.

*Ae. detritus* overwinter as larvae and the eggs can remain dormant and viable for long periods (Enevoldson, 2025). After the exclusion of the 2017 and 2018 data and applying the common window analysis, there was a positive association between delayed seasonal activity and the magnitude of the spring peak the following year. Such positive demographic responses to phenological shifts have been observed in other multivoltine species (Macgregor et al., 2019). The extension of autumn activity may lengthen the window for oviposition by female mosquitoes, with benefits in reproductive output once temperatures rise again the following spring. We note this analysis was not robust to every sampling window control, but it does provide an intriguing hypothesis to test, given that many temperate mosquito species over-winter with consequences for transmission (Folly et al., 2022).

An extended period of adult activity into autumn could increase the risk of arbovirus transmission in the UK. A longer vector activity season increases both the time available for viral amplification within mosquito populations and the temporal overlap with avian hosts – a mechanism recently identified as a key risk factor for WNV establishment in Britain (Ewing et al., 2021; Kirby et al., 2025). Research on UK arbovirus vectors has focused on *Culex pipiens*, but the detection of West Nile virus RNA in *Ae. vexans* in 2023 indicates that transmission risk in some settings may not be confined to a single mosquito species (Bruce et al., 2025), and vector competence and host-feeding behaviour can both vary among species and among populations of the same species (Kilpatrick et al., 2010, 2006, 2005). *Aedes detritus* is a competent West Nile virus vector under laboratory conditions (Blagrove et al., 2016), and its high-marsh habitat brings it into contact with both passerines and wildfowl, although contemporary data on its host-feeding preferences are lacking. Whether a longer period of late-season activity translates into greater contact with late-migrating birds, and therefore greater amplification opportunity, is an open question that our data cannot resolve directly, but it identifies a specific, testable link between phenological change and disease risk in this system.

Taken together, our findings indicate that a specialist coastal mosquito can show measurable phenological change over a relatively short run of years, expressed mainly as an extension of late season activity, and that this change is best captured by long-term, full-season monitoring rather than shorter or fixed-window surveys. As the risk of zoonotic mosquito-borne viruses establishing in the UK continues to be assessed, sustained investment in weekly, full-season monitoring of coastal vector species, beyond the timescale of individual funding cycles, would be necessary to detect and interpret such changes with confidence.

## Declarations

### Author contributions

**Nadja C. Wipf**: conceptualization, data curation, visualization, methodology, writing– original draft, writing–review and editing; **Thyge P. Enevoldson**: investigation, data curation, resources, writing–review and editing; **Christopher M. Jones**: conceptualization, funding acquisition, project administration, writing–original draft, writing–review and editing; **Jennifer S. Lord**: methodology, funding acquisition, project administration, writing–review and editing; **Tijani A. Sulaimon**: conceptualization, data curation, formal analysis, methodology, visualization, writing–original draft, writing–review and editing.

## Acknowledgements

We thank the residents of Neston who allowed us to place mosquito traps in their gardens.

## Data availability statement

The code and data used to produce this manuscript are available from the GitHub repository: https://github.com/NadjaWipf/saltmarsh-mosquito-model

## References

1. Bartoń, K., 2026. MuMIn: Multi-Model Inference.

2. Bell, J.R., Alderson, L., Izera, D., Kruger, T., Parker, S., Pickup, J., Shortall, C.R., Taylor, M.S., Verrier, P., Harrington, R., 2015. Long-term phenological trends, species accumulation rates, aphid traits and climate: five decades of change in migrating aphids. Journal of Animal Ecology 84, 21–34. 10.1111/1365-2656.12282

3. Bell, J.R., Botham, M.S., Henrys, P.A., Leech, D.I., Pearce-Higgins, J.W., Shortall, C.R., Brereton, T.M., Pickup, J., Thackeray, S.J., 2019. Spatial and habitat variation in aphid, butterfly, moth and bird phenologies over the last half century. Global Change Biology 25, 1982–1994. 10.1111/gcb.14592

4. Blagrove, M.S.C., Sherlock, K., Chapman, G.E., Impoinvil, D.E., McCall, P.J., Medlock, J.M., Lycett, G., Solomon, T., Baylis, M., 2016. Evaluation of the vector competence of a native UK mosquito Ochlerotatus detritus (Aedes detritus) for dengue, chikungunya and West Nile viruses. Parasites Vectors 9, 452. 10.1186/s13071-016-1739-3

5. Bloomfield, J.P., Bowler, D.E., Gripenberg, S., Harvey, M.C., 2026. Citizen science data reveals that earlier and shorter flight periods correlate with climate warming in the dark-edged bee-fly. Insect Conservation and Diversity. 10.1111/icad.70114

6. Bradley, T.J., Strange, K., Phillips, J.E., 1984. Osmotic and Ionic Regulation in Saline-Water Mosquito Larvae, in: Pequeux, A., Gilles, R., Bolis, L. (Eds.), Osmoregulation in Estuarine and Marine Animals: Proceedings of the Invited Lectures to a Symposium Organized within the 5th Conference of the European Society for Comparative Physiology and Biochemistry - Taormina, Sicily, Italy, September 5–8, 1983. Springer, Berlin, Heidelberg, pp. 35–50. 10.1007/978-3-642-45574-2_3

7. Bruce, R.C., Abbott, A.J., Jones, B.P., Gardner, B.L., Gonzalez, E., Ionescu, A.-M., Jagdev, M., Jenkins, A., Santos, M., Seilern-Macpherson, K., Hanmer, H.J., Robinson, R.A., Vaux, A.G., Johnson, N., Cunningham, A.A., Lawson, B., Medlock, J.M., Folly, A.J., 2025. Detection of West Nile virus via retrospective mosquito arbovirus surveillance, United Kingdom, 2025. Euro Surveill 30, 2500401. 10.2807/1560-7917.ES.2025.30.28.2500401

8. Burnham, K.P., Anderson, D.R., 2002. Model selection and multimodel inference: a practical information-theoretic approach. Springer, New York, NY.

9. Chandler, P., 2025. Checklist of Diptera of the British Isles [WWW Document]. Checklist of Diptera of the British Isles. URL https://dipterists.org.uk/checklist (accessed 12.15.25).

10. Chen, I.-C., Hill, J.K., Ohlemüller, R., Roy, D.B., Thomas, C.D., 2011. Rapid Range Shifts of Species Associated with High Levels of Climate Warming. Science 333, 1024–1026. 10.1126/science.1206432

11. Clarkson, M.J., Enevoldson, T.P., 2021. The factors which influence the breeding and number of Aedes detritus in the Neston area of Cheshire, UK, the production of a local mosquito forecast and public bite reporting. Journal of the European Mosquito Control Association 38, 17–32.

12. Clarkson, M.J., Setzkorn, C., 2011. The domestic mosquitoes of the Neston area of Cheshire, UK. European Mosquito Bulletin 29, 122–128.

13. Cooke, R., Outhwaite, C.L., Bladon, A.J., Millard, J., Rodger, J.G., Dong, Z., Dyer, E.E., Edney, S., Murphy, J.F., Dicks, L.V., Hui, C., Jones, J.I., Newbold, T., Purvis, A., Roy, H.E., Woodcock, B.A., Isaac, N.J.B., 2025. Integrating multiple evidence streams to understand insect biodiversity change. Science 388, eadq2110. 10.1126/science.adq2110

14. Didham, R.K., Basset, Y., Collins, C.M., Leather, S.R., Littlewood, N.A., Menz, M.H.M., Müller, J., Packer, L., Saunders, M.E., Schönrogge, K., Stewart, A.J.A., Yanoviak, S.P., Hassall, C., 2020. Interpreting insect declines: seven challenges and a way forward. Insect Conservation and Diversity 13, 103–114. 10.1111/icad.12408

15. Enevoldson, T.P., 2025. Long term viability of Aedes detritus (Haliday) and Aedes caspius (Pallas) (Diptera, Culicidae) eggs. Dipterists Digest 32, 36–38.

16. Ewing, D.A., Purse, B.V., Cobbold, C.A., White, S.M., 2021. A novel approach for predicting risk of vector-borne disease establishment in marginal temperate environments under climate change: West Nile virus in the UK. J. R. Soc. Interface. 18, 20210049. 10.1098/rsif.2021.0049

17. Folly, A.J., Sewgobind, S., Hernández-Triana, L.M., Mansfield, K.L., Lean, F.Z.X., Lawson, B., Seilern-Moy, K., Cunningham, A.A., Spiro, S., Wrigglesworth, E., Pearce-Kelly, P., Herdman, T., Johnston, C., Berrell, M., Vaux, A.G.C., Medlock, J.M., Johnson, N., 2022. Evidence for overwintering and autochthonous transmission of Usutu virus to wild birds following its redetection in the United Kingdom. Transboundary and Emerging Diseases 69, 3684–3692. 10.1111/tbed.14738

18. Forrest, J.R., 2016. Complex responses of insect phenology to climate change. Current Opinion in Insect Science, Global change biology * Molecular physiology 17, 49–54. 10.1016/j.cois.2016.07.002

19. Giraud, T., Hernangómez, D., Hijmans, R.J., Graham, H.A., McBain, M., 2025. maptiles: Download and Display Map Tiles.

20. Glazaczow, A., Orwin, D., Bogdziewicz, M., 2016. Increased temperature delays the late-season phenology of multivoltine insect. Sci Rep 6, 38022. 10.1038/srep38022

21. Guralnick, R.P., Campbell, L.P., Belitz, M.W., 2023. Weather anomalies more important than climate means in driving insect phenology. Commun Biol 6, 490. 10.1038/s42003-023-04873-4

22. Hartig, F., Lohse, L., leite, M. de S., 2024. DHARMa: Residual Diagnostics for Hierarchical (Multi-Level / Mixed) Regression Models.

23. Hawkes, F., Medlock, J.M., Vaux, A.G.C., Cheke, R., Gibson, G., 2020. Wetland mosquito survey handbook: assessing suitability of British wetlands for mosquitoes. Natural Resources Institute, Chatham, UK.

24. Hijmans, R.J., Brown, A., Barbosa, M., Dyba, K., Bivand, R., Chirico, M., Cordano, E., Pebesma, E., Rowlingson, B., Sumner, M.D., 2026. terra: Spatial Data Analysis.

25. Hill, J.K., Griffiths, H.M., Thomas, C.D., 2011. Climate Change and Evolutionary Adaptations at Species’ Range Margins. Annual Review of Entomology 56, 143–159. 10.1146/annurev-ento-120709-144746

26. Jones, C.M., Reynolds, D.R., Wipf, N.C., Chapman, J.W., Lehmann, T., 2025. The migratory behaviour of salt marsh mosquitoes: Revisiting the evidence. Medical and Veterinary Entomology n/a. 10.1111/mve.70021

27. Kilpatrick, A.M., Daszak, P., Jones, M.J., Marra, P.P., Kramer, L.D., 2006. Host heterogeneity dominates West Nile virus transmission. Proc Biol Sci 273, 2327–2333. 10.1098/rspb.2006.3575

28. Kilpatrick, A.M., Fonseca, D.M., Ebel, G.D., Reddy, M.R., Kramer, L.D., 2010. Spatial and Temporal Variation in Vector Competence of Culex pipiens and Cx. restuans Mosquitoes for West Nile Virus. Am J Trop Med Hyg 83, 607–613. 10.4269/ajtmh.2010.10-0005

29. Kilpatrick, A.M., Kramer, L.D., Campbell, S.R., Alleyne, E.O., Dobson, A.P., Daszak, P., 2005. West Nile Virus Risk Assessment and the Bridge Vector Paradigm. Emerg Infect Dis 11, 425–429. 10.3201/eid1103.040364

30. Kirby, G., Brown, R.E., Lee, M., Parvy, J.-P., Krabbendam, S., Pondeville, E., Johnston, C., Medlock, J.M., Vaux, A.G.C., Nelli, L., Baldini, F., Ferguson, H.M., 2026. Ecology of mosquitoes in Scottish wetlands and confirmation of Culex pipiens biotype molestus and pipiens/molestus hybrids in Scotland. Parasites Vectors. 10.1186/s13071-026-07474-2

31. Kirby, G., Vaux, A.G.C., Ferguson, H.M., Medlock, J.M., 2025. Ecological risk factors for the establishment of West Nile virus in Britain. Trends in Parasitology 41, 138–149. 10.1016/j.pt.2024.12.003

32. Macgregor, C.J., Thomas, C.D., Roy, D.B., Beaumont, M.A., Bell, J.R., Brereton, T., Bridle, J.R., Dytham, C., Fox, R., Gotthard, K., Hoffmann, A.A., Martin, G., Middlebrook, I., Nylin, S., Platts, P.J., Rasteiro, R., Saccheri, I.J., Villoutreix, R., Wheat, C.W., Hill, J.K., 2019. Climate-induced phenology shifts linked to range expansions in species with multiple reproductive cycles per year. Nat Commun 10, 4455. 10.1038/s41467-019-12479-w

33. Mackenzie-Impoinvil, L., Impoinvil, D.E., Galbraith, S.E., Dillon, R.J., Ranson, H., Johnson, N., Fooks, A.R., Solomon, T., Baylis, M., 2015. Evaluation of a temperate climate mosquito, Ochlerotatus detritus (=Aedes detritus), as a potential vector of Japanese encephalitis virus. Medical and Veterinary Entomology 29, 1–9. 10.1111/mve.12083

34. Marshall, J.F., 1938. The British Mosquitoes. British Museum (Natural History), London.

35. Marshall, J. F., 1933. An Inland Record of Aëdes detritus, Haliday (Diptera, Culicidæ). Nature 132, 135–135. 10.1038/132135a0

36. Met Office, Hollis, D., Carlisle, E., Kendon, M., Packman, S., Doherty, A., 2026. HadUK-Grid Gridded Climate Observations on a 1km grid over the UK, v1.3.2.ceda (1836-2025). 10.5285/789B3065D74A4C948AB05D33556C86D0

37. Moritz, S., Gatscha, S., Wang, E., Hause, R., 2025. imputeTS: Time Series Missing Value Imputation.

38. Rippel, T.M., Tomasula, J., Murphy, S.M., Wimp, G.M., 2021. Global change in marine coastal habitats impacts insect populations and communities. Current Opinion in Insect Science, Global change biology. Molecular physiology. October (2021) 47, 1–6. 10.1016/j.cois.2021.02.010

39. Roy, D.B., Oliver, T.H., Botham, M.S., Beckmann, B., Brereton, T., Dennis, R.L.H., Harrower, C., Phillimore, A.B., Thomas, J.A., 2015. Similarities in butterfly emergence dates among populations suggest local adaptation to climate. Glob Chang Biol 21, 3313–3322. 10.1111/gcb.12920

40. Sanders, C.J., Shortall, C.R., England, M., Harrington, R., Purse, B., Burgin, L., Carpenter, S., Gubbins, S., 2019. Long-term shifts in the seasonal abundance of adult Culicoides biting midges and their impact on potential arbovirus outbreaks. Journal of Applied Ecology 56, 1649–1660. 10.1111/1365-2664.13415

41. Schmidt, T.L., Endersby-Harshman, N., Mills, T., Rane, R., Pandey, G., Hardy, C., Court, L., Webb, C., Trewin, B., Neilan, B., Hoffmann, A.A., 2025. Populations of the Australian Saltmarsh Mosquito Aedes vigilax Vary Between Panmixia and Temporally Stable Local Genetic Structure. Evolutionary Applications 18, e70119. 10.1111/eva.70119

42. Service, M.W., 1972. An inland freshwater population of the salt marsh mosquito, Aedes detritus (Haliday). Journal of Entomology Series A, General Entomology 46, 117–121. 10.1111/j.1365-3032.1972.tb00117.x

43. Service, M.W., 1968. The ecology of the immature stages of Aedes detritus (Diptera: Culicidae). Journal of Applied Ecology 5, 613–630. 10.2307/2401636

44. Shortall, C.R., Cook, S.M., Mauchline, A.L., Bell, J.R., 2024. Long-term trends in migrating Brassicogethes aeneus in the UK. Pest Management Science 80, 2294–2305. 10.1002/ps.7501

45. Shortall, C.R., Moore, A., Smith, E., Hall, M.J., Woiwod, I.P., Harrington, R., 2009. Long-term changes in the abundance of flying insects. Insect Conservation and Diversity 2, 251–260. 10.1111/j.1752-4598.2009.00062.x

46. Tegar, S., Brass, D.P., Purse, B.V., Cobbold, C.A., White, S.M., 2026. Temperature-sensitive incubation, transmissibility and risk of Aedes albopictus-borne chikungunya virus in Europe. J. R. Soc. Interface. 23, 20250707. 10.1098/rsif.2025.0707

47. Wagner, D.L., 2020. Insect Declines in the Anthropocene. Annu Rev Entomol 65, 457–480. 10.1146/annurev-ento-011019-025151

48. Wood, S., 2025. mgcv: Mixed GAM Computation Vehicle with Automatic Smoothness Estimation.

49. Wood, S.N., 2017. Generalized additive models: An introduction with R, 2nd ed. Chapman and Hall / CRC Press.

50. Yang, G.-J., Brook, B.W., Bradshaw, C.J.A., 2009. Predicting the Timing and Magnitude of Tropical Mosquito Population Peaks for Maximizing Control Efficiency. PLOS Neglected Tropical Diseases 3, e385. 10.1371/journal.pntd.0000385

