## supplementary_material for "Environmental drivers of abundance and later seasonal activity of a salt marsh mosquito"

### Supporting information

*Supplementary figures*

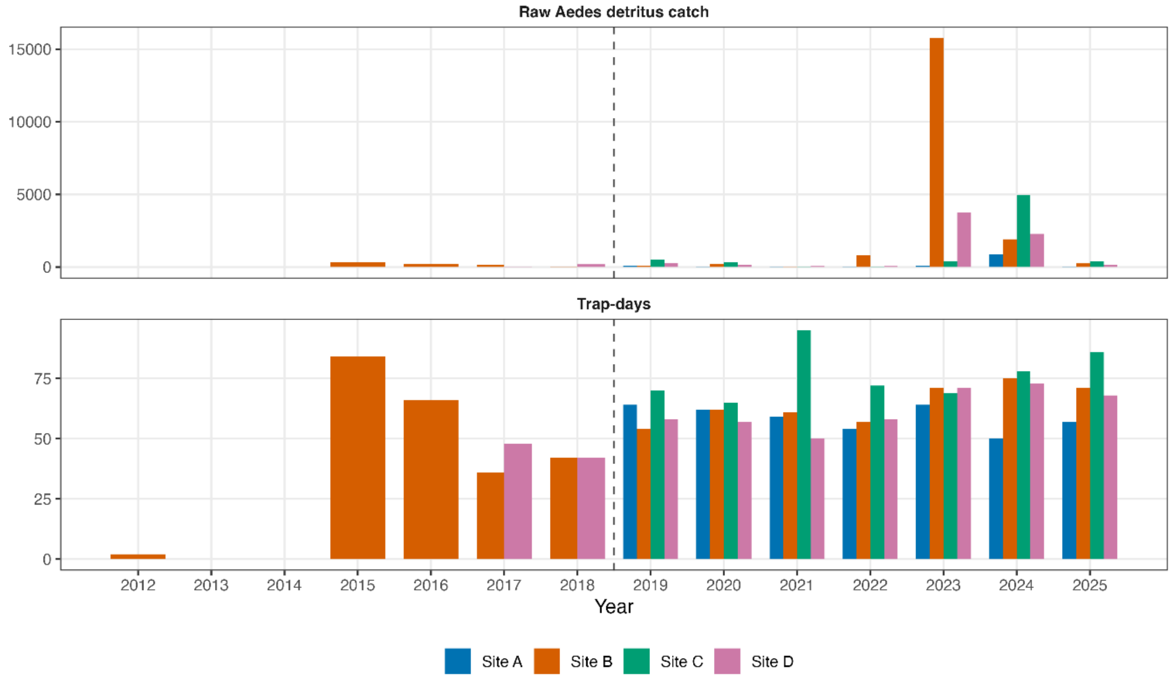


**Figure S1**. Raw *Aedes detritus* catch and trap-days at the four main sites by year. The dashed line marks the start of the consistent four-site trapping period used for monthly abundance modelling.

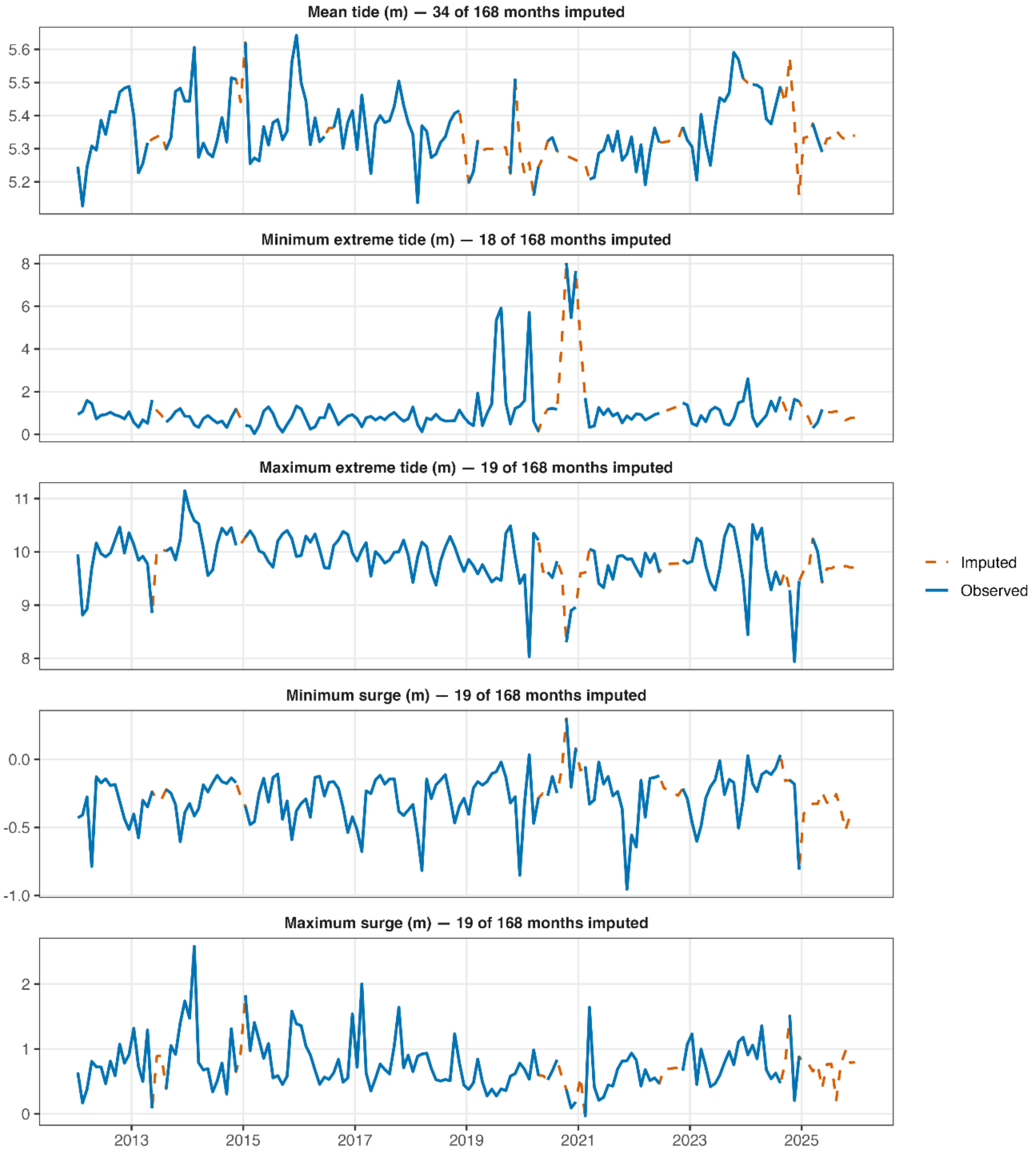


**Figure S2.** Observed and completed BODC monthly tidal series. Points show directly observed monthly summaries; connecting lines include values completed by the specified time-series imputation.

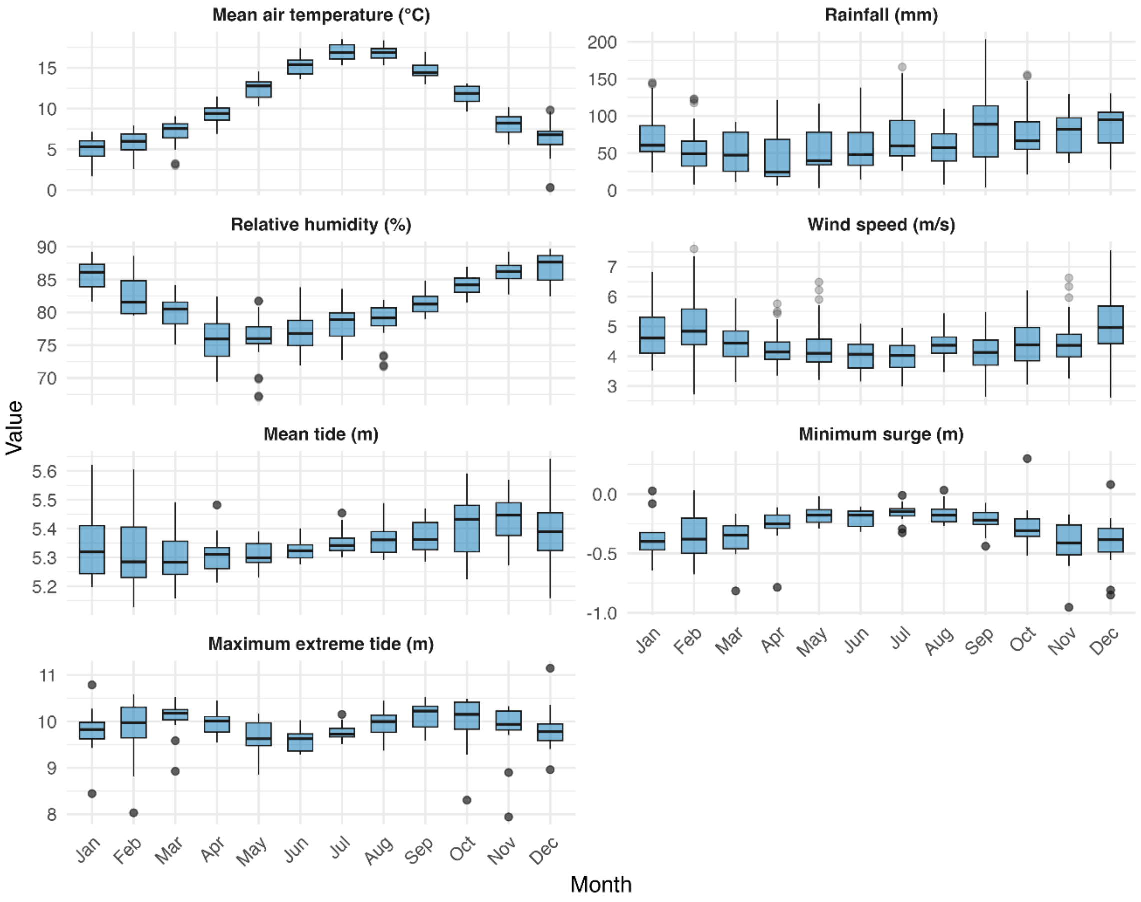


**Figure S3.** Monthly distributions of climate and tidal covariates used to examine seasonality and select the two candidate representations of seasonal structure.

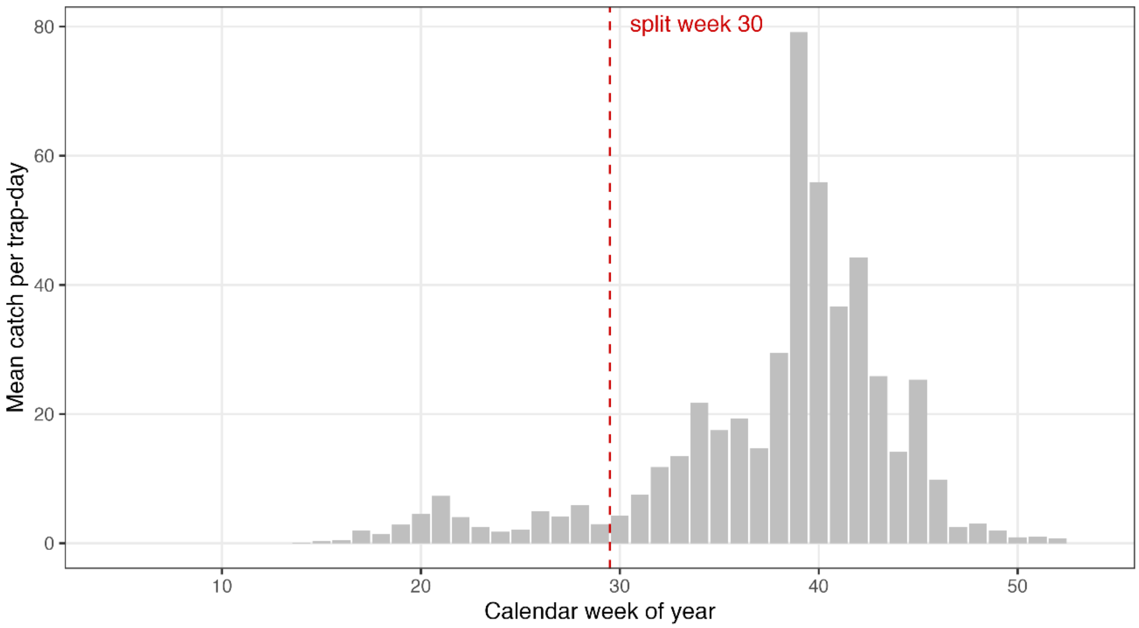


**Figure S4.** Mean weekly activity climatology. Calendar week 30 is the first week after the observed mid-season trough and separates the early- and late-season peak windows.

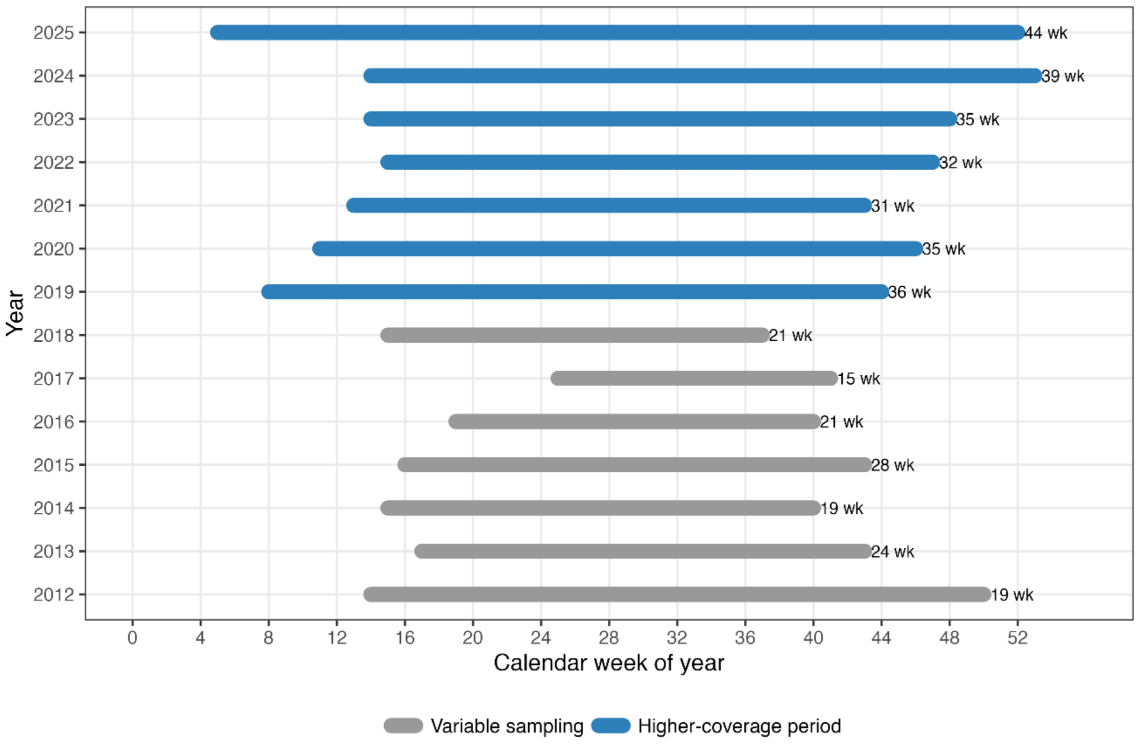


**Figure S5.** First and last sampled calendar weeks by year, with the number of sampled weeks. The late start in 2017 and early stop in 2018 motivated the metric-specific boundary controls.

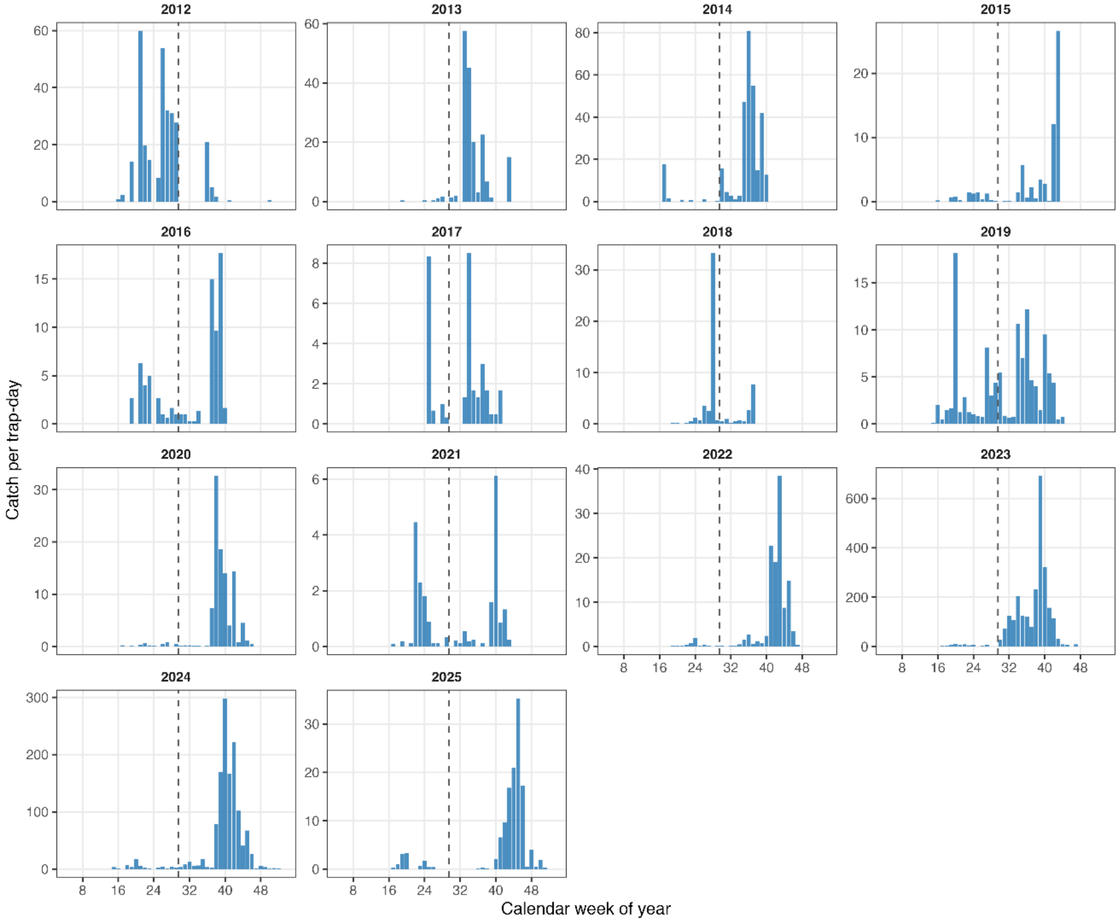


**Figure S6**. Weekly catch rate by year, pooled across trapping sites. The panels show year-to-year variation in the early-season cohort, the mid-season trough and the more consistent late-season cohort, together with gaps in sampling coverage.

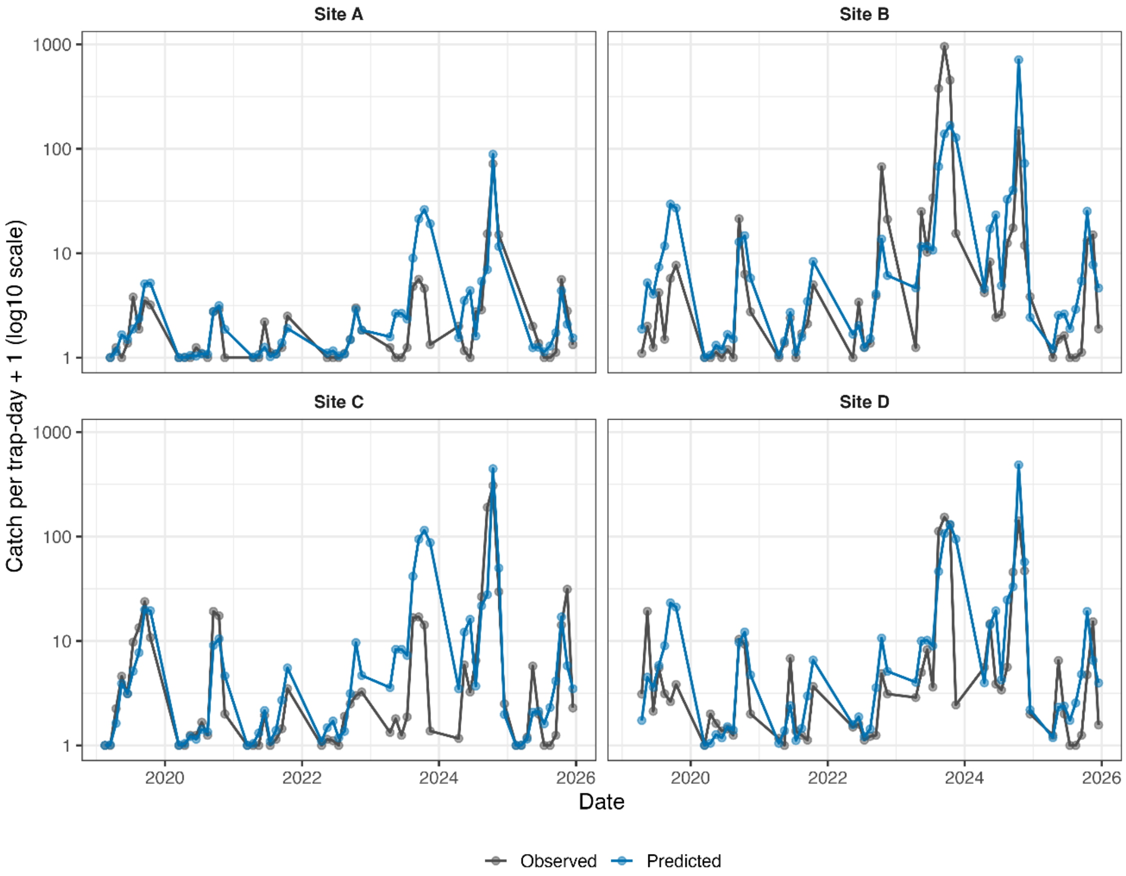


**Figure S7.** Observed and fitted monthly catch per trap-day by site for the selected negative-binomial GAM. This is an in-sample assessment of model fit, not an evaluation of out-of-sample prediction.

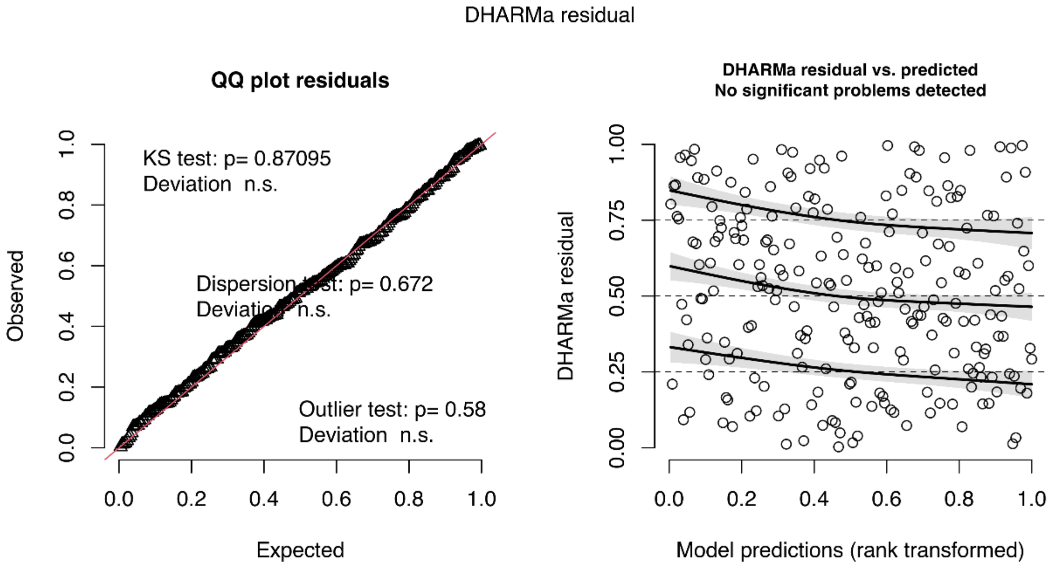


**Figure S8.** Simulation-based residual diagnostics for the selected monthly negative-binomial GAM. The panels show the DHARMa quantile-quantile check and scaled residuals against fitted values.

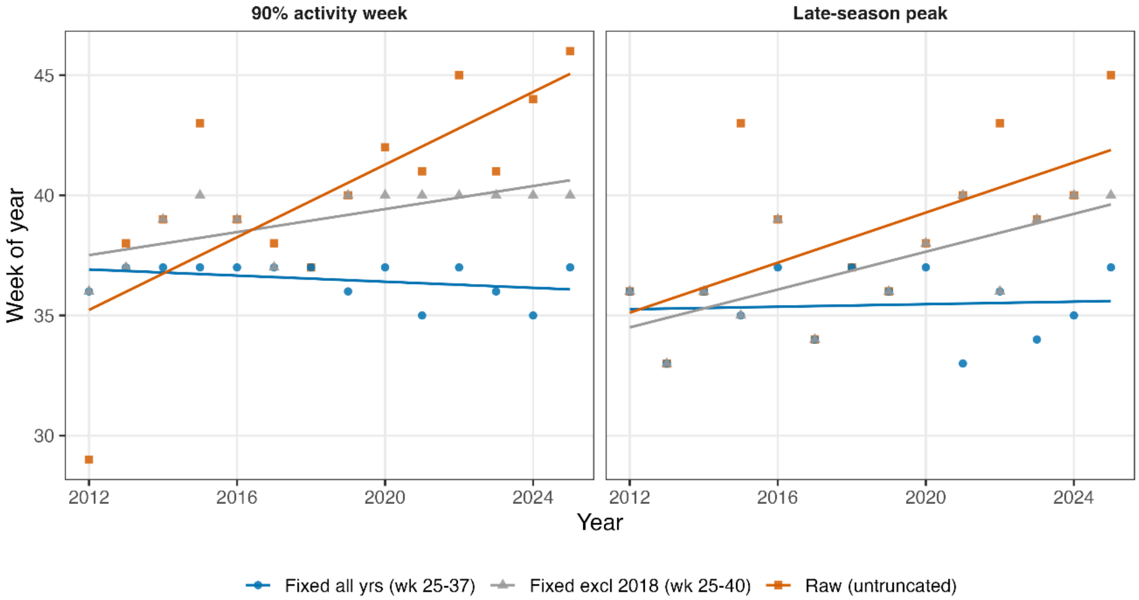

**Figure S9.** Trends in the 90% activity week and late-season peak under the raw untruncated data, the all-year common boundary window (weeks 25–37), and the metric-specific common boundary window excluding the early-stopping year 2018 (weeks 25–40). Points show annual metrics and lines show OLS trends.

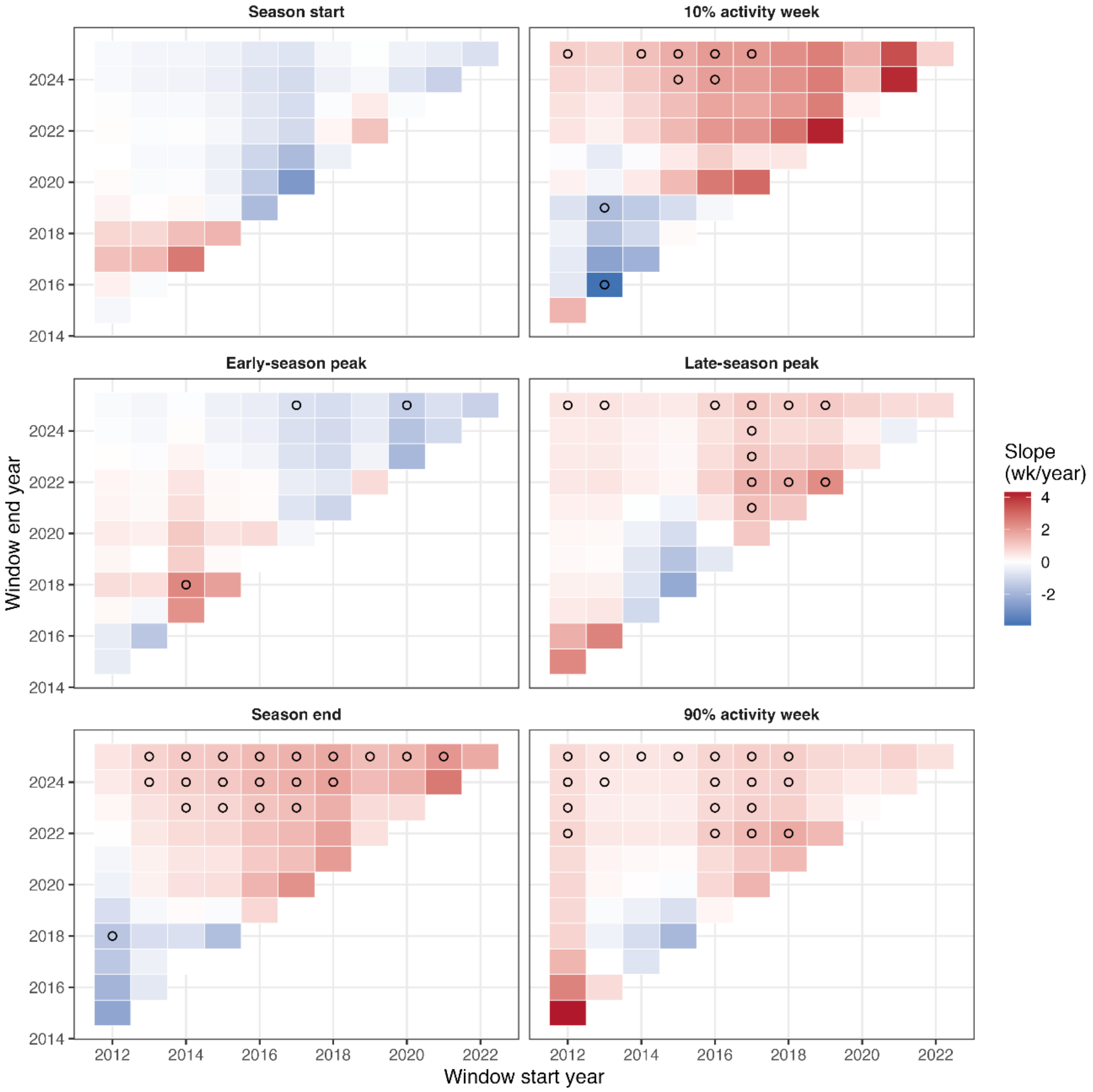


**Figure S10.** Phenology slopes across every contiguous period of at least four years. Each cell is one start-end year window, colour shows the fitted OLS slope, and open circles denote nominal $p < 0.05$.

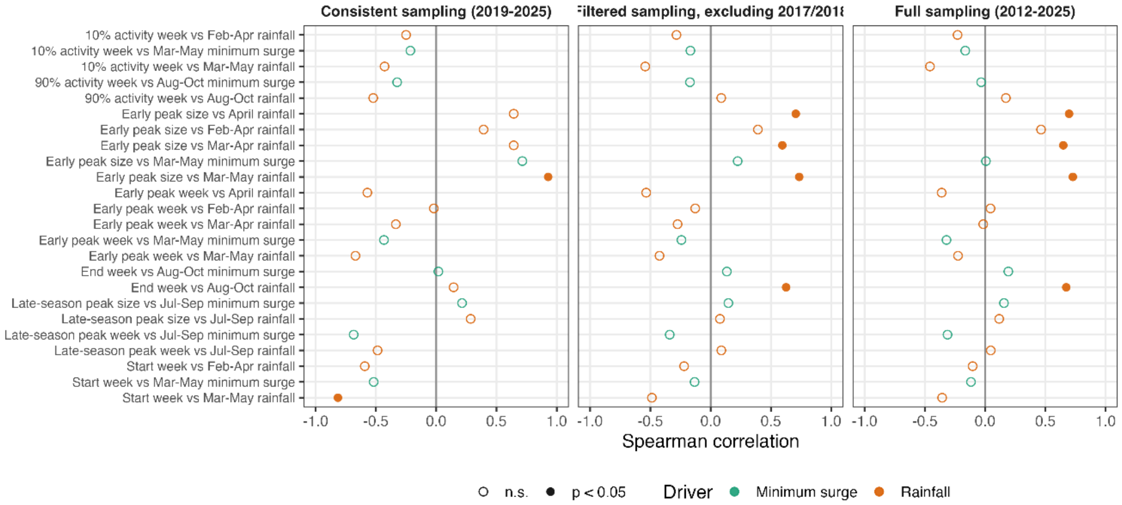


**Figure S11.** Complete exploratory annual phenology-driver correlations for the consistent sampling period (2019–2025), the full series excluding 2017 and 2018, and the full series (2012–2025). Orange points denote rainfall and green points minimum surge; filled points denote nominal $p < 0.05$. Values are given in Tables S10–S12.

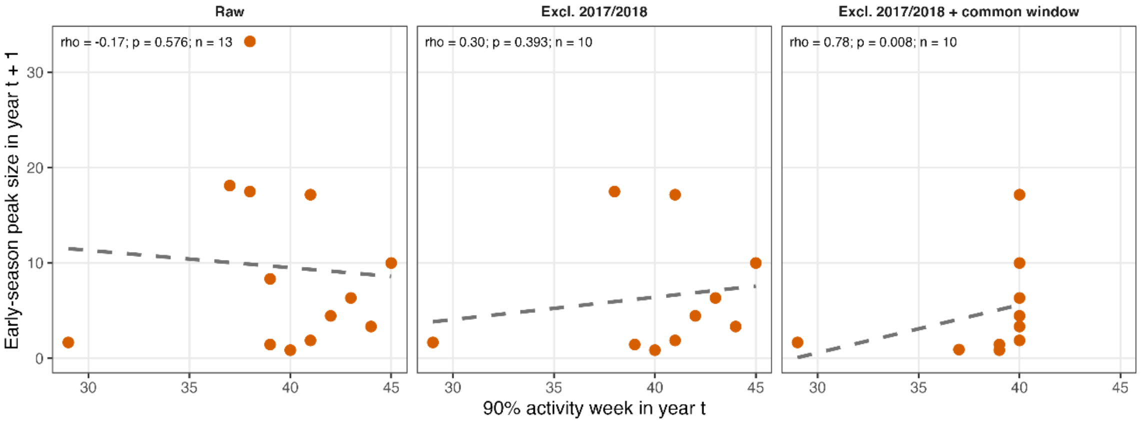


**Figure S12.** Association between the 90% activity week in year $t$ and early-season peak magnitude in year $t + 1$ under the raw data, after excluding 2017/2018, and after additionally applying the common-window control. Dashed OLS lines are visual guides; the annotations report Spearman correlations.

*Supplementary tables*

**Table S1.** Smooth terms in the selected monthly GAM.

| Term | edf | Reference df | Chi-square | *p*-value |
|---|---|---|---|---|
| Season | 6.78 | 7.63 | 166.70 | <0.001 |
| Site | 2.88 | 3 | 64.14 | <0.001 |
| Lagged rainfall | 1 | 1 | 37.27 | <0.001 |
| Minimum surge | 1 | 1 | 14.11 | <0.001 |

*Note*: Selected model: S + R + Su. Estimated degrees of freedom (edf) indicate the complexity of each smooth effect.

**Table S2.** Model-selection sensitivity excluding 2023.

| Hypothesis | Model | df | AICc | ΔAICc | weight |
|---|---|---|---|---|---|
| H1 | S + R + Su + W | 22 | 1388.2 | 0.0 | 0.90 |
| H1 | S + R + Su | 19 | 1394.1 | 5.9 | 0.05 |
| H1 | S + R + W | 21 | 1395.2 | 7.1 | 0.03 |
| H1 | S + R + Xt + W | 25 | 1395.2 | 7.1 | 0.03 |
| H1 | S + R + Xt | 22 | 1400.4 | 12.2 | 0 |
| H1 | S + R | 18 | 1401.1 | 12.9 | 0 |
| H2 | T + H + R + Su | 20 | 1420.8 | 32.6 | 0 |
| H2 | T + H + R + Su + W | 22 | 1422.9 | 34.7 | 0 |
| H2 | T + H + R | 18 | 1429.9 | 41.7 | 0 |
| H1 | S | 17 | 1430.1 | 41.9 | 0 |
| H2 | T + H + R + Xt | 19 | 1431.1 | 42.9 | 0 |
| H2 | T + H + R + W | 21 | 1432.7 | 44.6 | 0 |
| H2 | T + H + R + Xt + W | 22 | 1434.2 | 46.0 | 0 |
| H2 | T + H | 17 | 1455.4 | 67.2 | 0 |
| Null | Null (site + year) | 9 | 1566.4 | 178.2 | 0 |
| Null | Null (intercept) | 2 | 1642.5 | 254.4 | 0 |

*Note*: Models are ranked by small-sample corrected Akaike Information Criterion (AICc). ΔAICc indicates the difference from the best model, and Akaike weights represent the relative probability that a given model is the best in the candidate set. S denotes the calendar-month smooth (seasonality), R one-month lagged rainfall (mm), Su minimum surge (m), W wind speed (m/s), Xt maximum extreme tide (m), T air temperature (°C), and H relative humidity (%). H1 are hypothesis with integrated seasonality; H2 with explicit environmental drivers; Null the null hypotheses. All models include site random effects, year as categorical factor, and a log trap-effort offset.

**Table S3**. Model-selection sensitivity using observed minimum surge. The candidate set was refitted to 177 site-months (77.3% of the abundance data) with directly observed minimum surge. The minimum-surge smooth in S + R + Su had $p$ <0.001.

| Hypothesis | Model | df | AICc | ΔAICc | weight |
|---|---|---|---|---|---|
| H1 | S + R + Su | 20 | 1402.0 | 0.0 | 0.52 |
| H1 | S + R + Su + W | 22 | 1402.3 | 0.3 | 0.44 |
| H1 | S + R + Xt | 21 | 1409.2 | 7.2 | 0.01 |
| H1 | S + R + Xt + W | 24 | 1409.7 | 7.7 | 0.01 |
| H2 | T + H + R + Xt | 20 | 1411.5 | 9.5 | 0 |
| H2 | T + H + R + Su | 20 | 1411.6 | 9.6 | 0 |
| H2 | T + H + R + Su + W | 21 | 1413.6 | 11.6 | 0 |
| H2 | T + H + R + Xt + W | 22 | 1414.0 | 12.0 | 0 |
| H1 | S + R + W | 21 | 1415.9 | 14.0 | 0 |
| H1 | S + R | 18 | 1417.3 | 15.3 | 0 |
| H2 | T + H + R | 18 | 1434.0 | 32.1 | 0 |
| H2 | T + H + R + W | 20 | 1436.5 | 34.5 | 0 |
| H1 | S | 17 | 1440.2 | 38.2 | 0 |
| H2 | T + H | 17 | 1449.1 | 47.1 | 0 |
| Null | Null (site + year) | 10 | 1532.6 | 130.6 | 0 |
| Null | Null (intercept) | 2 | 1634.5 | 232.6 | 0 |

*Note*: Models are ranked by small-sample corrected Akaike Information Criterion (AICc). ΔAICc indicates the difference from the best model, and Akaike weights represent the relative probability that a given model is the best in the candidate set. S denotes the calendar-month smooth (seasonality), R one-month lagged rainfall (mm), Su minimum surge (m), W wind speed (m/s), Xt maximum extreme tide (m), T air temperature (°C), and H relative humidity (%). H1 are hypothesis with integrated seasonality; H2 with explicit environmental drivers; Null the null hypotheses. All models include site random effects, year as categorical factor, and a log trap-effort offset.

**Table S4**. Annual phenology trend estimates.

| Period | Metric | Years | OLS slope | 95% CI | OLS *p* |
|---|---|---|---|---|---|
| 2019-2025 | Season start | 7 | -0.036 | [-0.90, 0.82] | 0.919 |
| 2019-2025 | 10% activity week | 7 | 2.571 | [-0.73, 5.87] | 0.101 |
| 2019-2025 | 90% activity week | 7 | 0.786 | [-0.03, 1.60] | 0.056 |
| 2019-2025 | Early-season peak | 7 | -0.571 | [-1.87, 0.73] | 0.309 |
| 2019-2025 | Late-season peak | 7 | 1.071 | [0.04, 2.11] | 0.045 |
| 2019-2025 | Season end | 7 | 1.321 | [0.40, 2.24] | 0.014 |
| 2012-2025 | Season start | 14 | -0.149 | [-0.54, 0.24] | 0.417 |
| 2012-2025 | 10% activity week | 14 | 0.965 | [0.07, 1.86] | 0.038 |
| 2012-2025 | 90% activity week | 14 | 0.756 | [0.34, 1.18] | 0.002 |
| 2012-2025 | Early-season peak | 14 | -0.141 | [-0.64, 0.36] | 0.553 |
| 2012-2025 | Late-season peak | 14 | 0.521 | [0.11, 0.93] | 0.018 |
| 2012-2025 | Season end | 14 | 0.527 | [-0.06, 1.11] | 0.073 |
| 2012-2025, excl. 2017/2018 | Season start | 12 | -0.092 | [-0.33, 0.15] | 0.410 |
| 2012-2025, excl. 2017/2018 | 10% activity week | 12 | 0.935 | [-0.06, 1.93] | 0.062 |
| 2012-2025, excl. 2017/2018 | 90% activity week | 12 | 0.740 | [0.29, 1.19] | 0.004 |
| 2012-2025, excl. 2017/2018 | Early-season peak | 12 | -0.108 | [-0.59, 0.38] | 0.629 |
| 2012-2025, excl. 2017/2018 | Late-season peak | 12 | 0.490 | [0.07, 0.91] | 0.027 |
| 2012-2025, excl. 2017/2018 | Season end | 12 | 0.485 | [-0.04, 1.01] | 0.068 |

*Note*: OLS slopes are shown with 95% confidence intervals and *p*-values.

**Table S5.** Complete robustness ladder for phenology trends.

| Metric | 2019-2025 | 2012-2025 | 2012-2025 excl. 17/18 | Fixed all | Fixed excl. 17/18 | Moving windows |
|---|---|---|---|---|---|---|
| Season start | -0.04 | -0.15 | -0.09 | -0.04 (wk 25-37) | -0.04 (wk 19-40) | 0/66 |
| Early-season peak | -0.57 | -0.14 | -0.11 | -0.03 (wk 25-37) | -0.30 (wk 19-40) | 3/66 (+/-) |
| 10% activity week | +2.57 | +0.96* | +0.93 | -0.15 (wk 25-37) | -0.05 (wk 19-40) | 9/66 (+/-) |
| Late-season peak | +1.07* | +0.52* | +0.49* | +0.03 (wk 25-37) | +0.38* (wk 19-40) | 12/66 (+) |
| 90% activity week | +0.79 | +0.76* | +0.74* | -0.06 (wk 25-37) | +0.43* (wk 19-40) | 19/66 (+) |
| Season end | +1.32* | +0.53 | +0.49 | +0.00 (wk 25-37) | +0.08* (wk 19-40) | 20/66 (+/-) |

*Note*: This audit table retains the boundary-sensitive first and last catch metrics and the 10% activity week.

**Table S6**. Exact fixed-window phenology trend estimates.

| Period | Window | Metric | Years | OLS slope | 95% CI | OLS p |
|---|---|---|---|---|---|---|
| Fixed 2019-2025 | wk 15-43 | Season start | 7 | 0.036 | [-0.71, 0.78] | 0.906 |
| Fixed 2019-2025 | wk 15-43 | 10% activity week | 7 | 0.429 | [-3.96, 4.82] | 0.812 |
| Fixed 2019-2025 | wk 15-43 | Early-season peak | 7 | -0.571 | [-1.87, 0.73] | 0.309 |
| Fixed 2019-2025 | wk 15-43 | Late-season peak | 7 | 0.857 | [-0.07, 1.79] | 0.064 |
| Fixed 2019-2025 | wk 15-43 | Season end | 7 | 0 | [0.00, 0.00] | NA |
| Fixed 2019-2025 | wk 15-43 | 90% activity week | 7 | 0.321 | [-0.14, 0.78] | 0.134 |
| Fixed 2012-2025 | wk 25-37 | Season start | 14 | -0.037 | [-0.10, 0.02] | 0.197 |
| Fixed 2012-2025 | wk 25-37 | 10% activity week | 14 | -0.149 | [-0.55, 0.25] | 0.431 |
| Fixed 2012-2025 | wk 25-37 | Early-season peak | 14 | -0.031 | [-0.19, 0.13] | 0.690 |
| Fixed 2012-2025 | wk 25-37 | Late-season peak | 14 | 0.026 | [-0.19, 0.24] | 0.796 |
| Fixed 2012-2025 | wk 25-37 | Season end | 14 | 0 | [0.00, 0.00] | NA |
| Fixed 2012-2025 | wk 25-37 | 90% activity week | 14 | -0.064 | [-0.17, 0.04] | 0.218 |
| Fixed 2012-2025, excl. 2017/2018 | wk 19-40 | Season start | 12 | -0.042 | [-0.13, 0.04] | 0.302 |
| Fixed 2012-2025, excl. 2017/2018 | wk 19-40 | 10% activity week | 12 | -0.055 | [-1.08, 0.97] | 0.907 |
| Fixed 2012-2025, excl. 2017/2018 | wk 19-40 | Early-season peak | 12 | -0.295 | [-0.70, 0.11] | 0.137 |
| Fixed 2012-2025, excl. 2017/2018 | wk 19-40 | Late-season peak | 12 | 0.375 | [0.13, 0.62] | 0.007 |
| Fixed 2012-2025, excl. 2017/2018 | wk 19-40 | Season end | 12 | 0.085 | [0.01, 0.16] | 0.033 |
| Fixed 2012-2025, excl. 2017/2018 | wk 19-40 | 90% activity week | 12 | 0.433 | [0.05, 0.82] | 0.031 |

*Note*: These estimates underlie the fixed-window columns in Table 2. Restricting all years to their shared weeks can remove genuine early- or late-season activity.

**Table S7**. Onset sensitivity excluding 2017.

| Period | Metric | Years | OLS slope | OLS p | Window |
|---|---|---|---|---|---|
| Raw, excl. 2017 only | Season start | 13 | -0.098 | 0.392 | untruncated |
| Raw, excl. 2017 only | 10% activity week | 13 | 0.944 | 0.050 | untruncated |
| Raw, excl. 2017 only | Early-season peak | 13 | -0.125 | 0.610 | untruncated |
| Fixed, excl. 2017 only | Season start | 13 | -0.041 | 0.286 | wk 19-37 |
| Fixed, excl. 2017 only | 10% activity week | 13 | -0.385 | 0.234 | wk 19-37 |
| Fixed, excl. 2017 only | Early-season peak | 13 | -0.309 | 0.151 | wk 19-37 |

*Note*: The late-start year 2017 is excluded to assess potential bias in season-start metrics.

**Table S8**. Late-season sensitivity excluding 2018.

| Period | Metric | Years | OLS slope | OLS p | Window |
|---|---|---|---|---|---|
| Raw, excl. 2018 only | Late-season peak | 13 | 0.518 | 0.023 | untruncated |
| Raw, excl. 2018 only | 90% activity week | 13 | 0.749 | 0.003 | untruncated |
| Raw, excl. 2018 only | Season end | 13 | 0.51 | 0.053 | untruncated |
| Fixed, excl. 2018 only | Late-season peak | 13 | 0.394 | 0.006 | wk 25-40 |
| Fixed, excl. 2018 only | 90% activity week | 13 | 0.24 | 0.005 | wk 25-40 |
| Fixed, excl. 2018 only | Season end | 13 | 0.082 | 0.032 | wk 25-40 |

*Note*: The early-stop year 2018 is excluded to assess potential bias in season-end metrics.

**Table S9.** Moving-window trend summary.

| Metric | All windows | Nominal $p < 0.05$, positive | Nominal $p < 0.05$, negative |
|---|---|---|---|
| Season end | 66 | 19 | 1 |
| 90% activity week | 66 | 19 | 0 |
| Late-season peak | 66 | 12 | 0 |
| 10% activity week | 66 | 7 | 2 |
| Early-season peak | 66 | 1 | 2 |
| Season start | 66 | 0 | 0 |

*Notes*: Windows overlap and are not independent. Counts are descriptive stability checks; nominal *p*-values are not multiple-testing-adjusted.

**Table S10**. Complete phenology-driver correlations for the full sampling period (2012–2025).

| Driver family | Metric | Years | Spearman rho | *p*-value | BH *q*-value |
|---|---|---|---|---|---|
| Rainfall | Start week vs Mar-May rainfall | 14 | -0.357 | 0.210 | 0.631 |
| Rainfall | 10% activity week vs Mar-May rainfall | 14 | -0.458 | 0.100 | 0.400 |
| Rainfall | Early peak week vs Mar-May rainfall | 14 | -0.225 | 0.439 | 0.879 |
| Rainfall | Early peak size vs Mar-May rainfall | 14 | 0.727 | 0.003 | 0.067 |
| Rainfall | Start week vs Feb-Apr rainfall | 14 | -0.104 | 0.724 | 0.914 |
| Rainfall | 10% activity week vs Feb-Apr rainfall | 14 | -0.229 | 0.431 | 0.879 |
| Rainfall | Early peak week vs Feb-Apr rainfall | 14 | 0.045 | 0.880 | 0.982 |
| Rainfall | Early peak size vs Feb-Apr rainfall | 14 | 0.464 | 0.095 | 0.400 |
| Rainfall | Early peak week vs Mar-Apr rainfall | 14 | -0.016 | 0.958 | 0.982 |
| Rainfall | Early peak size vs Mar-Apr rainfall | 14 | 0.648 | 0.012 | 0.073 |
| Rainfall | Early peak week vs April rainfall | 14 | -0.361 | 0.205 | 0.631 |
| Rainfall | Early peak size vs April rainfall | 14 | 0.697 | 0.006 | 0.067 |
| Rainfall | Late-season peak week vs Jul-Sep rainfall | 14 | 0.047 | 0.875 | 0.982 |
| Rainfall | Late-season peak size vs Jul-Sep rainfall | 14 | 0.116 | 0.692 | 0.914 |
| Rainfall | End week vs Aug-Oct rainfall | 14 | 0.673 | 0.008 | 0.067 |
| Rainfall | 90% activity week vs Aug-Oct rainfall | 14 | 0.172 | 0.557 | 0.891 |
| Minimum surge | Start week vs Mar-May minimum surge | 14 | -0.117 | 0.689 | 0.914 |
| Minimum surge | 10% activity week vs Mar-May minimum surge | 14 | -0.165 | 0.573 | 0.891 |
| Minimum surge | Early peak week vs Mar-May minimum surge | 14 | -0.321 | 0.264 | 0.665 |
| Minimum surge | Early peak size vs Mar-May minimum surge | 14 | 0.007 | 0.982 | 0.982 |
| Minimum surge | Late-season peak week vs Jul-Sep minimum surge | 14 | -0.312 | 0.277 | 0.665 |
| Minimum surge | Late-season peak size vs Jul-Sep minimum surge | 14 | 0.156 | 0.594 | 0.891 |
| Minimum surge | End week vs Aug-Oct minimum surge | 14 | 0.192 | 0.510 | 0.891 |
| Minimum surge | 90% activity week vs Aug-Oct minimum surge | 14 | -0.033 | 0.911 | 0.982 |

*Note*: February-April rainfall is lag-aligned and March-May rainfall represents the UK spring activity period. Benjamini-Hochberg (BH) q-values adjust all 24 rainfall and minimum-surge tests within this period.

**Table S11**. Complete phenology-driver correlations for the consistent sampling period (2019-2025).

| Driver family | Metric | Years | Spearman rho | p-value | BH q-value |
|---|---|---|---|---|---|
| Rainfall | Start week vs Mar-May rainfall | 7 | -0.815 | 0.025 | 0.305 |
| Rainfall | 10% activity week vs Mar-May rainfall | 7 | -0.429 | 0.337 | 0.578 |
| Rainfall | Early peak week vs Mar-May rainfall | 7 | -0.670 | 0.100 | 0.409 |
| Rainfall | Early peak size vs Mar-May rainfall | 7 | 0.929 | 0.003 | 0.060 |
| Rainfall | Start week vs Feb-Apr rainfall | 7 | -0.593 | 0.161 | 0.481 |
| Rainfall | 10% activity week vs Feb-Apr rainfall | 7 | -0.250 | 0.589 | 0.737 |
| Rainfall | Early peak week vs Feb-Apr rainfall | 7 | -0.020 | 0.967 | 0.969 |
| Rainfall | Early peak size vs Feb-Apr rainfall | 7 | 0.393 | 0.383 | 0.613 |
| Rainfall | Early peak week vs Mar-Apr rainfall | 7 | -0.335 | 0.463 | 0.675 |
| Rainfall | Early peak size vs Mar-Apr rainfall | 7 | 0.643 | 0.119 | 0.409 |
| Rainfall | Early peak week vs April rainfall | 7 | -0.571 | 0.180 | 0.481 |
| Rainfall | Early peak size vs April rainfall | 7 | 0.643 | 0.119 | 0.409 |
| Rainfall | Late-season peak week vs Jul-Sep rainfall | 7 | -0.487 | 0.268 | 0.536 |
| Rainfall | Late-season peak size vs Jul-Sep rainfall | 7 | 0.286 | 0.535 | 0.713 |
| Rainfall | End week vs Aug-Oct rainfall | 7 | 0.144 | 0.758 | 0.827 |
| Rainfall | 90% activity week vs Aug-Oct rainfall | 7 | -0.523 | 0.229 | 0.508 |
| Minimum surge | Start week vs Mar-May minimum surge | 7 | -0.519 | 0.233 | 0.508 |
| Minimum surge | 10% activity week vs Mar-May minimum surge | 7 | -0.214 | 0.645 | 0.737 |
| Minimum surge | Early peak week vs Mar-May minimum surge | 7 | -0.433 | 0.331 | 0.578 |
| Minimum surge | Early peak size vs Mar-May minimum surge | 7 | 0.714 | 0.071 | 0.409 |
| Minimum surge | Late-season peak week vs Jul-Sep minimum surge | 7 | -0.685 | 0.090 | 0.409 |
| Minimum surge | Late-season peak size vs Jul-Sep minimum surge | 7 | 0.214 | 0.645 | 0.737 |
| Minimum surge | End week vs Aug-Oct minimum surge | 7 | 0.018 | 0.969 | 0.969 |
| Minimum surge | 90% activity week vs Aug-Oct minimum surge | 7 | -0.324 | 0.478 | 0.675 |

*Note*: February-April rainfall is lag-aligned and March-May rainfall represents the UK spring activity period. Benjamini-Hochberg (BH) q-values adjust all 24 rainfall and minimum-surge tests within this period.

**Table S12**. Complete phenology-driver correlations for the full sampling period excluding 2017 and 2018.

| Driver family | Metric | Years | Spearman rho | *p*-value | BH *q*-value |
|---|---|---|---|---|---|
| Rainfall | Start week vs Mar-May rainfall | 12 | -0.488 | 0.108 | 0.370 |
| Rainfall | 10% activity week vs Mar-May rainfall | 12 | -0.543 | 0.068 | 0.293 |
| Rainfall | Early peak week vs Mar-May rainfall | 12 | -0.424 | 0.169 | 0.508 |
| Rainfall | Early peak size vs Mar-May rainfall | 12 | 0.734 | 0.007 | 0.123 |
| Rainfall | Start week vs Feb-Apr rainfall | 12 | -0.220 | 0.491 | 0.786 |
| Rainfall | 10% activity week vs Feb-Apr rainfall | 12 | -0.284 | 0.372 | 0.776 |
| Rainfall | Early peak week vs Feb-Apr rainfall | 12 | -0.128 | 0.691 | 0.790 |
| Rainfall | Early peak size vs Feb-Apr rainfall | 12 | 0.392 | 0.208 | 0.555 |
| Rainfall | Early peak week vs Mar-Apr rainfall | 12 | -0.275 | 0.388 | 0.776 |
| Rainfall | Early peak size vs Mar-Apr rainfall | 12 | 0.594 | 0.042 | 0.249 |
| Rainfall | Early peak week vs April rainfall | 12 | -0.535 | 0.073 | 0.293 |
| Rainfall | Early peak size vs April rainfall | 12 | 0.706 | 0.010 | 0.123 |
| Rainfall | Late-season peak week vs Jul-Sep rainfall | 12 | 0.089 | 0.784 | 0.812 |
| Rainfall | Late-season peak size vs Jul-Sep rainfall | 12 | 0.077 | 0.812 | 0.812 |
| Rainfall | End week vs Aug-Oct rainfall | 12 | 0.625 | 0.030 | 0.237 |
| Rainfall | 90% activity week vs Aug-Oct rainfall | 12 | 0.088 | 0.786 | 0.812 |
| Minimum surge | Start week vs Mar-May minimum surge | 12 | -0.134 | 0.679 | 0.790 |
| Minimum surge | 10% activity week vs Mar-May minimum surge | 12 | -0.168 | 0.601 | 0.790 |
| Minimum surge | Early peak week vs Mar-May minimum surge | 12 | -0.242 | 0.448 | 0.786 |
| Minimum surge | Early peak size vs Mar-May minimum surge | 12 | 0.224 | 0.484 | 0.786 |
| Minimum surge | Late-season peak week vs Jul-Sep minimum surge | 12 | -0.340 | 0.280 | 0.671 |
| Minimum surge | Late-season peak size vs Jul-Sep minimum surge | 12 | 0.147 | 0.649 | 0.790 |
| Minimum surge | End week vs Aug-Oct minimum surge | 12 | 0.134 | 0.677 | 0.790 |
| Minimum surge | 90% activity week vs Aug-Oct minimum surge | 12 | -0.172 | 0.593 | 0.790 |

*Note*: February-April rainfall is lag-aligned and March-May rainfall represents the UK spring activity period. Benjamini-Hochberg (BH) q-values adjust all 24 rainfall and minimum-surge tests within this period.

**Table S13**. Exploratory consecutive-year carry-over correlations.

| Analysis | Metric | Years | Spearman rho | *p*-value |
|---|---|---|---|---|
| Raw all consecutive pairs | 90% activity week | 13 | -0.171 | 0.576 |
| Raw all consecutive pairs | Activity span (10-90%) | 13 | -0.207 | 0.498 |
| Filtered (excl. 2017/2018), no common window | 90% activity week | 10 | 0.304 | 0.393 |
| Filtered (excl. 2017/2018), no common window | Activity span (10-90%) | 10 | -0.544 | 0.104 |
| Filtered common window, excluding 2017/2018 | 90% activity week | 10 | 0.781 | 0.008 |
| Filtered common window, excluding 2017/2018 | Activity span (10-90%) | 10 | 0 | 1.000 |

*Not*e: These correlations test whether later or longer activity in year t is associated with a larger early-season peak in year $t + 1$
